# NASP maintains histone H3–H4 homeostasis through two distinct H3 binding modes

**DOI:** 10.1101/2021.11.02.467034

**Authors:** Hongyu Bao, Massimo Carraro, Valentin Flury, Yanhong Liu, Min Luo, Liu Chen, Anja Groth, Hongda Huang

## Abstract

Histone chaperones regulate all aspects of histone metabolism. NASP is a major histone chaperone for H3–H4 dimers critical for preventing histone degradation. Here, we identify two distinct histone binding modes of NASP and reveal how they cooperate to ensure histone H3–H4 supply. We determine the structures of a sNASP dimer, a complex of sNASP with an H3 α3 peptide, and the sNASP–H3–H4–ASF1b co-chaperone complex. This captures distinct functionalities of NASP and identifies two distinct binding modes involving the H3 α3 helix and the H3 αN region, respectively. Functional studies demonstrate the H3 αN-interaction represents the major binding mode of NASP in cells and shielding of the H3 αN region by NASP is essential in maintaining the H3–H4 histone soluble pool. In conclusion, our studies uncover the molecular basis of NASP as a major H3–H4 chaperone in guarding histone homeostasis.

## INTRODUCTION

Histone chaperones form a protein network that participates in all aspects of histone metabolism, including histone folding, posttranslational modification, transport, nucleosome assembly and disassembly, histone recycling and turnover, storage, and degradation (1–4). Thus, histone chaperones directly or indirectly influence genome stability and integrity, gene transcription, DNA replication and repair (5–7). Histone chaperones are a diverse class of proteins lacking sequence similarity that have evolved to shield histones in a myriad of ways (1). Structural studies of histones in complex with their chaperones have thus been critical to reveal the molecular basis of histone chaperone function and instrumental for functional studies. The structure of the DAXX–H3.3–H4 complex explained DAXX’s histone H3.3 variant specificity (8,9), and structure-guided functional study of TONSL revealed unique histone chaperone and reader functions in DNA replication and repair (10). More recently, structure-guided proteomics unraveled DNAJC9’s dual histone and heat-shock co-chaperone functions (11), linking ATP-resourced protein folding to nucleosome assembly pathways. Here, we focus on the structure of the nuclear autoantigenic sperm protein (NASP) and its interaction with diverse histone substrates. NASP is a conserved H3–H4 histone chaperone with central roles in histone metabolism, yet the molecular basis of sNASP escorting and safeguarding histone substrates remain elusive.

In human, NASP has two non-allelic splicing variants, the testicular NASP (tNASP) (12) and somatic NASP (sNASP) (13). tNASP is highly expressed in testes, ovarian and transformed cells, while sNASP is ubiquitously expressed. tNASP is suggested to function as a co-chaperone of HSP90 (14) and take part in the early folding of H3–H4 dimers (15), while sNASP has more broad roles in the chaperone network (15,16). sNASP is part of a multichaperone complex containing the histone chaperones ASF1 (a and b), RbAp46/48 and the histone acetyltransferase HAT1, involved in acetylation and transport of newly synthesized H3–H4 dimers (15). This multichaperone complex buffers new soluble H3–H4 dimers accumulating during replicational stress (17,18) and might also re-acetylate evicted H3–H4 dimers (16). Notably, sNASP and tNASP are uniquely required for maintaining the soluble pool of H3–H4 dimers during the cell cycle and protect them from degradation by chaperone-mediated autophagy (19). Recently, sNASP was also proposed to maintain a soluble pool of monomeric H3 in the nucleus (20). Consistent with its role as a major H3–H4 chaperone, homozygous deletion of *NASP* gene is embryonic lethal in mice (21).

NASP is broadly distributed across eukaryotes and its histone chaperone function is largely conserved (22). The fission yeast (*Schizosaccharomyces pombe*) homologue Sim3, the Arabidopsis (*Arabidopsis thaliana*) NASP and the budding yeast (*Saccharomyces cerevisiae*) homologue Hif1p were shown to bind and chaperone histones CENPA and H3 (with and without H4) (23–25). In addition, the frog (*Xenopus laevis*) homologue N1/N2 may function as a chaperone of stored maternal H3–H4 dimers in Xenopus oocytes (26,27). NASP and its homologues share an atypical tetratricopeptide repeat (TPR) domain. They constitute the SHNi-TPR (Sim3-Hif1-NASP interrupted TPR) family (23), predicted to have four TPR motifs with the TPR2 motif being interrupted by a long acidic region, a capping helix region immediately following the TPR4 motif, and a disordered C-terminal region containing a nuclear localization signal (NLS) (12,22,23,25,28,29). The recently determined structure of Hif1p mostly confirmed the structural features of the atypical SHNi-TPR domain (30). However, it has not been feasible to obtain structures of NASP family chaperones bound to histones H3 and H4 (31,32).

A variety of studies have biochemically characterized NASP and its interaction with diverse histone substrates *in vitro*. sNASP can form both a monomer and a dimer in solution (15,33), it can bind to histone H3 monomers and to H3–H4 dimers (34), and it can associate with ASF1 (a and b) to co-chaperone H3–H4 (15,35). Moreover, both sNASP and tNASP display nucleosome assembly activities toward canonical H3 and its variants (36,37). Recent work from the Ladurner lab revealed that sNASP can bind to an H3 C-terminal epitope via a central channel in its TPR domain (31), but that the chaperone uses a secondary interaction site(s) to bind H3–H4 in a co-chaperone complex with ASF1 (35). Here, we report the crystal structures of the sNASP dimer, the complex of sNASP with an H3 α3 peptide, and the sNASP–H3–H4–ASF1b co-chaperone complex. We identify two distinct H3 binding modes of NASP and our functional studies show that the H3 αN-binding mode represents the major binding mode in cells, adopted in co-chaperone complexes with ASF1 (a and b) and HAT1. Furthermore, we demonstrate that the shielding of the H3 αN region by NASP is essential in maintaining the H3–H4 histone soluble pool. In conclusion, our studies provide the molecular basis of NASP function as a major H3–H4 chaperone in maintaining histone homeostasis.

## MATERIAL AND METHODS

### Cloning and protein preparation in bacteria

The cDNA frangments of the human sNASP (amino acids, a.a. 1–340), sNASP (a.a. 30–340), sNASP (a.a. 30–340, with a deletion Δ101–159; hereafter referred to as ‘sNASP core’, sNASPc), sNASP (a.a. 1–340, with a deletion Δ101–159) and the budding yeast (Saccharomyces cerevisiae) full-length Hif1p were repectively cloned into a modified RSFDuet-1 vector (Novagen) with an N-terminal His_6_-SUMO tag. The cDNA fragments of the human full-length ASF1a, ASF1a (a.a. 1–155), sNASPc, H3.3 (a.a. 1–59), and the frog (*Xenopus laevis*) N1/N2 (a.a. 23–495, with a deletion Δ97–314; hereafter referred to as ‘N1/N2 core’, N1/N2c) were repectively cloned into the pGEX-6P-1 vector (GE Healthcare) resulting an N-terminal GST-tag. We used ClonExpress II One Step Cloning Kit (Vazyme, Nanjing) to generate these clones. All mutations were introduced using a standard PCR procedure and verified by DNA sequencing. For expression of these proteins, BL21 (DE3)-RIL (Stratagene) E. Coli cells were first transformed with the corresponding plasmids and then cultured using Luria-Bertani (LB) medium supplemented with proper antibiotics at 37 °C to an OD_600_ of 1.0-1.2. Then, protein expression was induced with 0.5 mM isopropyl α-D-1-thiogalactopyranoside (IPTG) and cells were further incubated overnight at 20 °C. For purification of the His_6_-SUMO tagged proteins, the expressed proteins were first purified by Ni Sepharose 6 Fast Flow beads (GE Healthcare). After removal of the His_6_-SUMO tag using home-made Ulp1 (SUMO protease), the proteins were further purified on a HiLoad 16/600 Superdex 200 column (GE Healthcare). For purification of the GST-tagged proteins, the expressed proteins were first purified by Glutathione Sepharose 4B beads (GE Healthcare), and then further purified with a HiLoad 16/600 Superdex 200 column (GE Healthcare). For some experiments, the GST-tags of GST-ASF1a (a.a. 1–155) and GST-H3.3 (a.a. 1–59) were removed with home-made 3C protease before the gel-filtration step.

For preparation of the sNASPc–H3 α3 complex, the sNASPc and H3.3 (a.a. 116–135) epitope were covalently linked into one expression cassette, and the cassette was cloned into the RSFDuet-1 vector with an N-terminal His_6_-SUMO tag. The expression and purification procedures were the same as described above. For preparation of the sNASPc–H3–H4–ASF1b heterotetramer: first, the sNASP (a.a. 30–323, Δ101–159) and ASF1b (a.a. 1–158) were covalently linked with a 8G-linker (GGGSGGGS) into one expression cassette, and the cassette was cloned into the RSFDuet-1 vector with an N-terminal His_6_-SUMO tag; then the resulting plasmid was coexpressed with a pETDuet-1 vector containing histones H3.3 and H4 genes. The resulting sNASPc–H3–H4–ASF1b heterotetramer was first purified by Ni Sepharose 6 Fast Flow beads (GE Healthcare). After removal of the His_6_-SUMO tag by home-made Ulp1 (SUMO protease), the complex was further purified by Heparin and HiLoad 16/600 Superdex 200 columns (GE Healthcare).

The preparation of H3.3–H4 and H3.3 (I51A R52A Y54A)–H4 dimers followed a similar procedure in our previous study (38) with small modifications. The histone complexes were first captured by a heparin column, and then further purified on a HiLoad 16/600 Superdex 200 column (GE Healthcare) with a high-salt buffer (20 mM Tris, pH 7.5, 2.0 M NaCl).

### Isothermal titration calorimetry (ITC)

ITC experiments were carried out on a MicroCal PEAQ-ITC (Malvern Panalytical Ltd) at 25 °C. The peptides except for the H3.3 (a.a. 1–59) peptide were all purchased from GenScript (Nanjing). The H3.3 (a.a. 1–59) peptide was expressed in E. coli and purified as described above. The buffer for our ITC experiments was 50 mM Tris, pH 7.5, 200 mM NaCl. For ITC assays between the H3 α3 (a.a. 116–135) peptide and different sNASP constructs, the titration protocol consisted of 19 injections of 0.4 mM H3 α3 peptide (in syringe) into 0.04 mM sNASP protein (in cell and counted as sNASP monomer). For ITC assays between the H3 N-terminal peptides, including the H3.3 (a.a. 1–59), H3.3 (a.a. 1–15), H3.3 (a.a. 16–39), H3.3 αN (a.a. 40–59), H3.3 αN R52A and H3.3 αN Y54A peptides, and different sNASP constructs, the titration protocol consisted of 19 injections of 0.8 mM peptide (in syringe) into 0.06 mM sNASP protein (in cell and counted as sNASP monomer). The data were processed with Microcal Origin software and the curves were fit to the ‘one set of sites’ model. ITC assays between the budding yeast Hifp and H3 peptides were performed in the same way.

### Crystallization

The purified sNASPc dimer at a concentration of 5 mg/ml was crystallized in 0.2 M Calcium acetate, 0.1 M HEPES, pH 7.5, 40% v/v PEG 400, with the sitting-drop vapor-diffusion method at 20 °C. The concentration of PEG 400 in the mother liquor was high enough to serve as cryoprotectant, thus all crystals were directly flash frozen in liquid nitrogen.

The sNASPc–H3 α3 complex at a concentration of 3 mg/ml was crystallized in 1.4 M Na-K phosphate, pH 8.2, with the sitting-drop vapor-diffusion method at 20 °C. The crystals were soaked in a cryoprotectant made form the mother liquor supplemented with 20% glycerol before being flash frozen in liquid nitrogen.

The sNASPc–H3–H4–ASF1b heterotetramer at a concentration of 10 mg/ml was crystallized in 8% v/v Tacsimate, pH 6.0, 20% w/v PEG 3,350, with the sitting-drop vapor-diffusion method at 20 °C. The crystals were soaked in a cryoprotectant made form the mother liquor supplemented with 10% glycerol before being flash frozen in liquid nitrogen.

### X-ray Data collection and structure determination

All X-ray diffraction data were collected on beamline 19U1 of the Shanghai Synchrotron Radiation Facility (SSRF) (39). The X-ray diffraction data of the sNASPc dimer, sNASPc–H3 α3 complex and sNASPc–H3–H4–ASF1b heterotetramer were collected at wavelengths of 0.9785 Å, 0.9766 Å and 0.9786 Å, respectively.

Data were processed and scaled using HKL3000 (40) and CCP4 program (41). The structure of the sNASPc–H3 α3 complex was solved by molecular replacement in PHASER (42) with an inital model derived from the crystal structure of the budding yeast Hif1p (PDB 4NQ0). After molecular replacement, the strucural model was manually built using Coot (43) and refined in PHENIX (44). The structure of the sNASPc dimer was solved by molecular replacement in PHASER with the structure of the sNASPc–H3 α3 complex as the search model, and were also manually built using Coot and refined in PHENIX. The structure of the sNASPc–H3–H4–ASF1b heterotetramer was solved by molecular replacement in PHASER with initial models derived from the structure of the sNASPc dimer and our previous structure of the MCM2–ASF1b–H3.3–H4 complex (PDB 5BNX), and were also manually built using Coot and refined in PHENIX. All the structural figures in this study were prepared with PyMOL (The PyMOL Molecular Graphics System, Schrödinger).

### GST pulldown assays

For pulldowns of the GST-ASF1a–H3–H4 or GST-ASF1a–H3(I51A R52A Y54A)–H4 complexes with sNASPc and its mutants: 50 μL of Glutathione Sepharose 4B beads was suspended with 200 μL of 1 M binding buffer (20 mM Tris, pH 7.5, 1.0 M NaCl); and 1.5 nmol of GST-tagged full length ASF1a and 2 nmol of H3.3–H4 dimer or H3.3(I51A R52A Y54A)–H4 dimer were added and incubated at 4 °C for 1 hour; then the beads were washed quickly once with 1 mL of 1 M washing buffer (1 M binding buffer, 0.5% v/v Triton X-100), and once with 1 mL of 1 M binding buffer, and twice with 1mL of 0.3 M binding buffer (20 mM Tris, pH 7.5, 0.3 M NaCl); then the beads were suspended with 200 μL of 0.3 M binding buffer, and 3 nmol of purified sNASPc or its mutants was added and incubated at 4 °C for 1 hour; finally, the beads were washed quickly four times with 1 mL of 0.3 M washing buffer (0.3 M binding buffer, 0.5% v/v Triton X-100) before addition of 50 μL of SDS-PAGE sample loading buffer.

For pulldowns of GST-sNASPc and its mutants with H3–H4 dimer: 50 μL of Glutathione Sepharose 4B beads was suspended with 200 μL of 0.3 M binding buffer; and 1.5 nmol of GST-sNASPc or its mutants was added and incubated at 4 °C for 20 min; then 3 nmol of purified H3.3–H4 dimer or H3.3 (I51A R52A Y54A)–H4 dimer was added and incubated at 4 °C for another 3 hours; then the beads were washed quickly four times with 1 mL of 0.75 M washing buffer (20 mM Tris, pH 7.5, 0.75 M NaCl, 0.5% v/v Triton X-100) before addition of 50 μL of SDS-PAGE sample loading buffer. Pulldowns of GST-N1/N2c and its mutants with H3.3–H4 dimer, and pulldowns of GST-sNASP^(30-340)^ and its mutants with H3.3–H4 dimer were performed in the same way. All the samples were analyzed with SDS-PAGE.

### Size-exclusion chromatography coupled to multi-angle light scattering (SEC-MALS)

For molar mass determination, the purified protein complexes were analyzed using an ÄKTA-MALS system at room temperature. For each experiment, 100 μL solution of 1.5 mg/mL protein was injected onto a Superdex 200 Increase 10/300 GL (GE Healthcare) column equilibrated in a buffer of 20 mM Tris, pH 7.5, 0.5 mM DTT and 0.5 M NaCl, at a flow rate of 0.5 mL per minute. Separation and ultraviolet detection were performed using an ÄKTA system (GE Healthcare). The ÄKTA system was coupled on-line to an 8-angle MALS detector (DAWN HELEOS II, Wyatt Technology) and a differential refractometer (Optilab T-rEX, Wyatt Technology). Data were analyzed using ASTRA 6.1.2.45. To detect the molar mass of the sNASPc–H3–H4–ASF1a heterotetramer, the purified sNASPc dimer was mixed with the H3–H4 dimer and ASF1a (a.a. 1–155) at a molar ratio of 1:2:2, which was then applied to the SEC-MALS assay. To detect the molar mass of the sNASPc–H3-H4 complex, the purified NASPc dimer was mixed with the H3–H4 dimer at a molar ratio of 1:2, which was then applied to the SEC-MALS assay.

### Antibodies

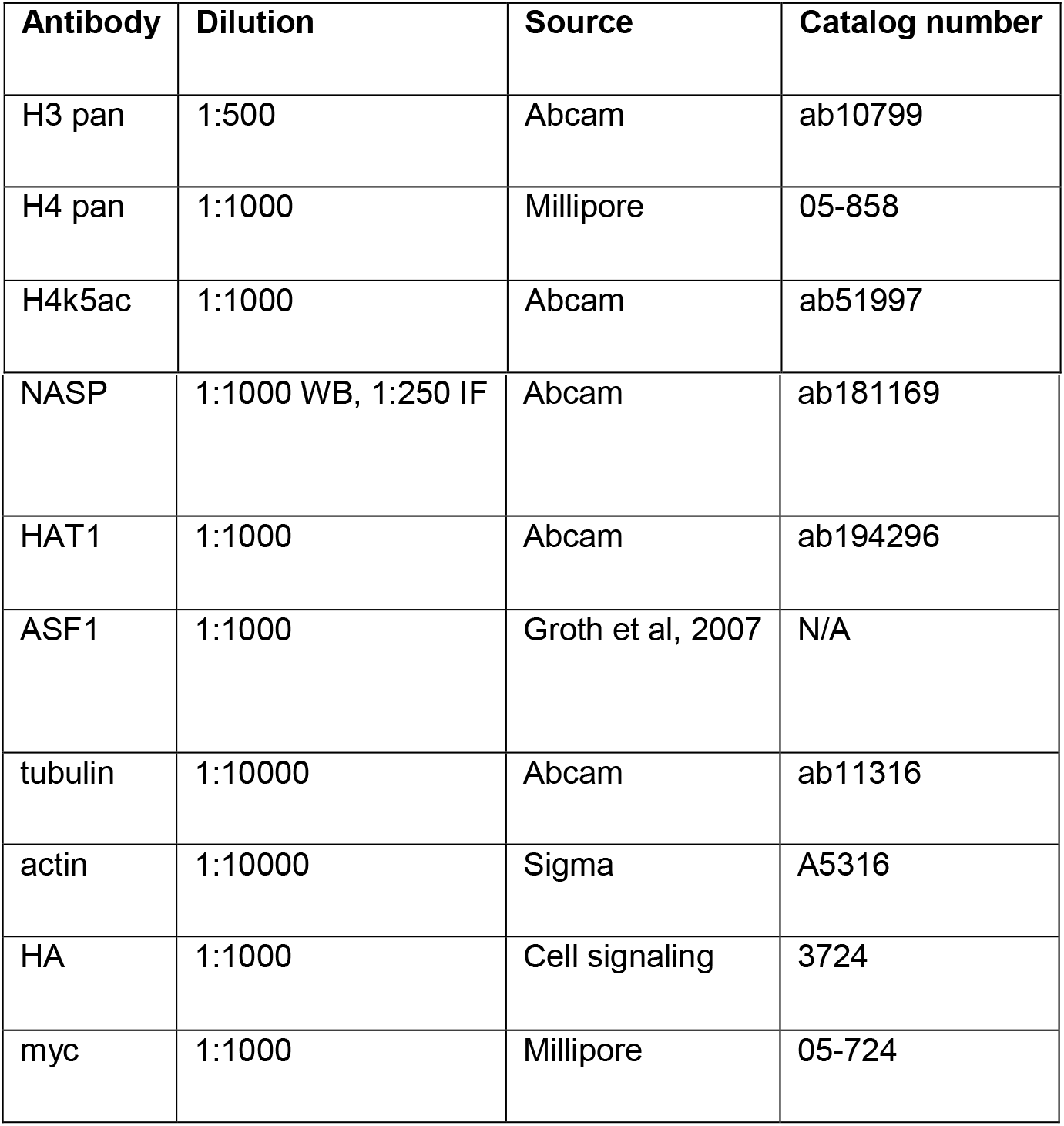

### Recombinant DNA

| Plasmid | Source | Reference |
| --- | --- | --- |
| pCMV6-sNASP-Myc-Flag (WT) | Origene | RC208783 |
| pCMV6-sNASP-V265E I272E-L282E-V286E-I300E-L307E-Myc-Flag (6E) | This study | N/A |
| pLVX-TetOne-Puro | Clontech | 631849 |
| pLVX-TetOne-Puro-TwinStrep-HA | This study | N/A |
| pLVX-TetOne-puro-sNASP-TwinStrep-HA (sNASP-Strep-HA WT) | This study | N/A |
| pLVX-TetOne-puro-sNASP-E177A-W180A-D181A-TwinStrep-HA (sNASP-Strep-HA EWD3A) | This study | N/A |
| pLVX-TetOne-puro-sNASP-E246A-Y249A-L253A-TwinStrep-HA (sNASP-Strep-HA EYL3A) | This study | N/A |
| pLVX-TetOne-puro-sNASP-E177A-W180A-D181A-E246A-Y249A-L253A-TwinStrep-HA (sNASP-Strep-HA EWD3A+EYL3A) | This study | N/A |
| pUC57-NASP-FKBP12<sup>F-36V</sup> | This study | N/A |
| pSpCas9(BB)-2A-Puro-V2.0 | Addgene | #62988 |
| pSpCas9(BB)-2A-Puro-NASP-sgRNA-V2.0 | This study | N/A |
| pVSV | Addgene | #138479 |
| psPAX2 | Addgene | #12260 |

### Primers

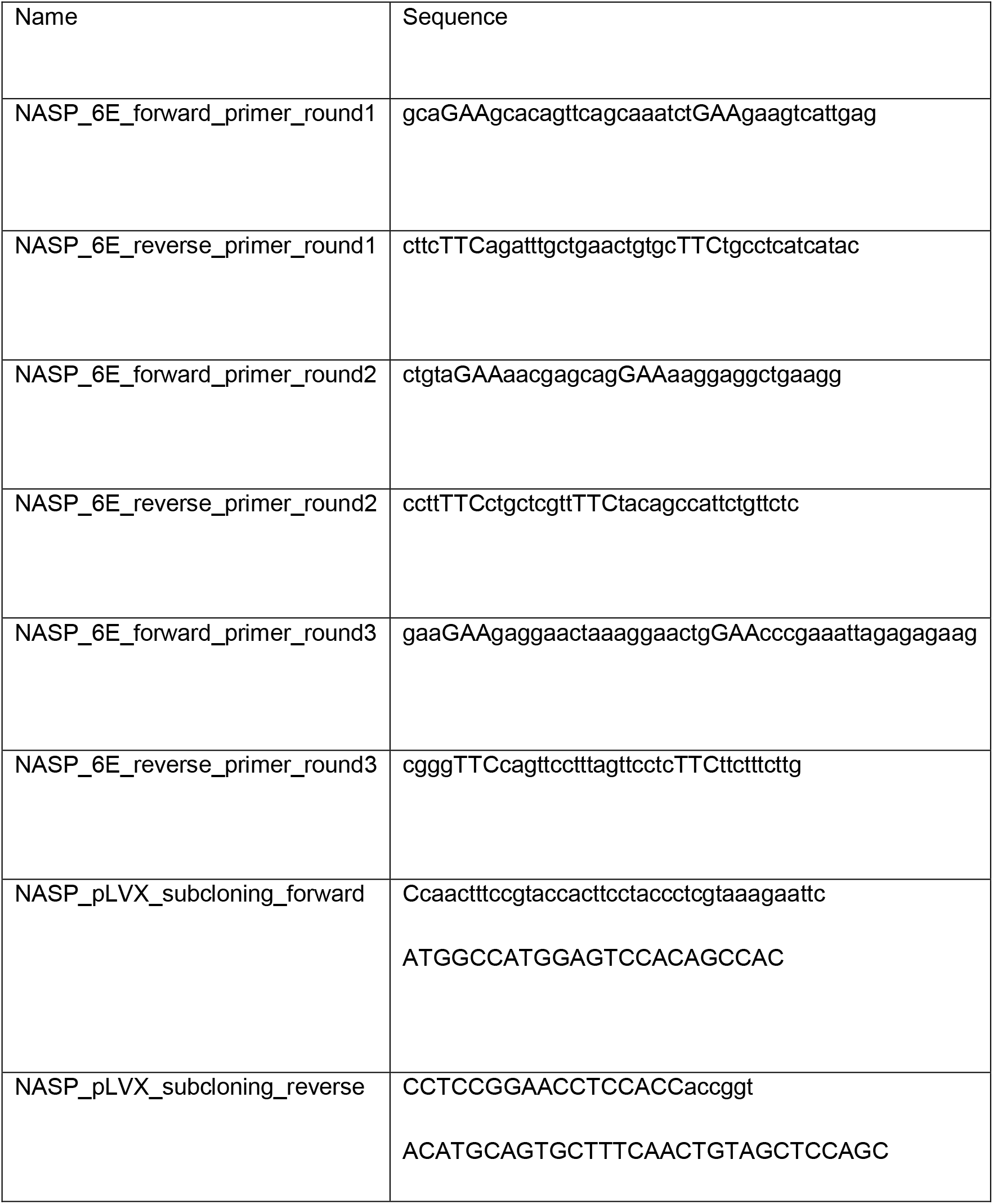

### Plasmid construction for cellular experiments

The sNASP dimerization mutant 6E (V265E I272E L282E V286E I300E L307E) was generated by three successive rounds of site directed mutagenesis of the pCMV6-sNASP-Myc-Flag vector. The C-terminal TwinStrep-HA tag (sequence TGGGGSGGGASWSHPQFEKGGSGGGSWSH PQFEKGGYPYDVPDYA*) was synthetized and cloned (by Genscript) between the EcoRI and BamHI sites of the pLVX-TetOne-Puro vector (631849, Clontech). Subsequently, sNASP coding sequence were sub-cloned into pLVX-TetOne-puro-TwinStrep-HA by amplifying sNASP cDNA with primers that also create the homologous arms and using these PCR product as “mega-primer” pair. NASP EWD3A(E177A-W180A-D181A), EYL3A (E246A-Y249A-L253A) and EWD3A+EYL3A (E177A-W180A-D181A-E246A-Y249A-L253A) mutants were synthetized and cloned (by Genscript) by site directed mutagenesis of pLVX-TetOne-puro-sNASP-TwinStrep-HA plasmid.

The pSpCas9(BB)-2A-Puro-NASP-sgRNA-V2.0, targeting the endogenous alleles of NASP in HCT116, was cloned between Bbsl sites of the pSpCas9(BB)-2A-Puro-V2.0 (62988, Addgene). The plasmid pUC57-NASP-FKBP12^F36V^, carrying homology arms for NASP alleles, was synthetized, and cloned by Genescript and used as a donor plasmid for genome editing.

Site directed mutagenesis was performed using established QuickChange mutagenesis protocols (Stratagene) or Infusion HD-directed mutagenesis (Takara). For Infusion HD-directed mutagenesis template plasmids were amplified with Phusion HF (F530S, Thermo Scientific) using mutagenic primers that also create homologous arms which, after PCR purification (28104, QIAgen) and Dpn1 digest (R0176L, NEB), were recombined through Infusion HD cloning (638933, Takara).

### Cell culture

Unless stated otherwise cells were grown in DMEM with Glutamax (35050061, Gibco), 10% FBS (12389802, Hyclone) and 1% penicillin/streptomycin (15140122, Gibco) at 37 °C with 5% CO_2_. HeLa S3 and HCT116 pLVX-TetOne-Puro cell lines were grown under Puromycin selection (1 μg/ml, P8833, Sigma). In Hela S3 and HCT116 pLVX-TetOne-Puro, the expression of sNASP-TwinStrep-HA, referred as sNASP-Strep-HA above, wt and mutants was induced by treatment with 2 μg/ml Doxycycline (631311, Clontech) for 24-48 hours. HCT116 NASP-FKBP12^F36V^ (NASP-dTAG) were treated with 0.2% DMSO (67-68-5, Sigma) or 5 μM dTAG-13 (6605, Tocris) for the indicated times, to induce the degradation of endogenous NASP. In the complementation experiments, HCT116 cells were simultaneously treated with 2 μg/ml Doxycycline and either 0.2% DMSO or 5 μM dTAG-13 for 48 hours, to ensure a synchronized degradation of the endogenous protein and expression of sNASP-Strep-HA wt or mutants. For cell viability experiments, HCT116 cells were seeded at low confluency in a 24 well plate. 24 hours after seeding, cells were treated with either 0.2% DMSO or 5 μM dTAG-13 for 48 hours, and cell viability was quantified with CellTiter-Blue® Cell Viability Assay (G8080, Promega), according to the manufacturer’s instructions.

### Genome Editing

In HCT116 cells, the C-terminal of NASP was fused in frame with FKBP12^F36V^ (dTAG), immediately before the stop codon. Cells were plated in a 6 well plate and co-transfected with 1 μg of pUC57-NASP-FKBP12^F36V^ (by Genscript) and 2 μg of pSpCas9(BB)-2A-Puro-NASP-sgRNA_V2.0 (#62988, Addgene) using Lipofectamine 3000 (L3000001, Thermo Fisher Scientific), according to the manufacturer’s instructions. 24 hours after transfection, cells were passaged to a 10 cm plate at the proper dilution to ensure single cell clones and selection media containing 1 μg/mL puromycin was added. 48 hours after the selection started, the selection media was removed and fresh media was added. From this point, single cell clones were isolated and moved in 96-well format. To isolate genomic DNA for genotyping, cells were washed twice with cold PBS, placed for 10 min at −80 °C and incubated with squishing buffer (10 mM Tris-HCl, pH 8; 1 mM EDTA, 25 mM NaCl, 200 μg/ml Proteinase K) at 65 °C for 1 hour. Proteinase K was inactivated by incubation at 95 °C for 10 minutes. 2 μl of this lysate was directly used for PCR reactions with 2× Taq start master mix (M0496L, NEB), and the two NASP clone screening primers (see primer table). Amplified DNA was sequenced using the same two primers.

### Lentivirus generation and transfection

Cell lines expressing sNASP wt and mutants from pLVX-TetOne-Puro constructs were created via lentiviral transduction of HCT116 or HeLa S3 suspension cells and Puromycin selection (1 μg/ml) 20-24 hours post-transduction. The lentivirus containing supernatants were harvested and filtered through a 0.45 μm syringe 40-60 hours after co-transfection of 293FT cells with 10 μg pLVX-TetOne-Puro, 5 μg pVSV (#138479, Addgene), 8 μg psPAX2 (#12260, Addgene) using Lipofectamine 2000 (11668019, Thermo Fisher Scientific) according to the manufacturer’s instructions. The presence of lentiviral particles was confirmed using Lenti-X GoStix Plus (631280, Clontech) according to the manufacturer’s instructions.

### Cell extracts

For whole cell extracts, cells were washed twice in cold PBS and lysed with Laemmli sample buffer 1X (LSB 4X: Tris-HcL (200 mM, pH 6.8), SDS 4%, glycerol (40%), DTT (100 mM)) supplemented with Benzonase (0.015 volumes, 25 U/μl, Millipore, 70746) and MgCl2 (0.01 volume, 1 M). Extracts were left one hour at 37 °C to digest DNA, boiled at 96 °C for 20 minutes and cleared by centrifugation (16,000 g, 10 mins).

For soluble extracts, cells were washed twice with cold PBS and pelleted by centrifugation (300 g, 3 mins) at 4 °C. The pellet was resuspended in ice cold NP40-NaCl buffer (300 mM NaCl, 0.05% Nonidet P40, 50 mM Tris-HCl pH 7.6, 0.1 mM EDTA, 5% glycerol) with freshly added inhibitors (NaF (5 mM) and β-Glycerolphosphate (10 mM), Phenylmethanesulfonyl fluoride (0.1 mM), Leupeptin (10 μg/ml), Pepstatin A (10 μg/ml), Trichostatin A (100 ng/ml), Na_3_VO_4_ (0.2 mM)) and left 15 minutes at 4 °C before the lysate was cleared by centrifugation (11,000 g, 20 mins), transferred to a new tube and cleared again (11,000 g, 10 mins) and used directly for immunoprecipitation experiments or stored at −80 °C.

### Immunoprecipitation and western blot analysis

Protein concentrations were measured using Pierce™ 660 nm Protein Assay Reagent (Thermo Scientific) or the Bradford protein assay (Bio-Rad). For immunoprecipitation, 2-3 mg of sNASP-FLAG-myc and sNASP-Strep-HA extracts were incubated for 3 hours at 4 °C degrees with 50 μl of anti-Flag M2 (A2220, Sigma) or Anti-TwinStrep Mag-Strep beads (2-4090-002, Iba). After incubation, the beads were washed five times in ice-cold wash buffer (150 mM NaCl, 0.02% Nonidet P40, 50 mM Tris-HCl pH 7.6, 0.1 mM EDTA, 5%glycerol). Bound proteins were eluted with either Laemmli sample buffer 1X (LSB 4X: Tris-HCl (200 mM, pH 6.8), SDS 4%, glycerol (40%), DTT (100 mM)) or biotin elution buffer 1X (BXT buffer, 2-1042-025, Iba), respectively. Western blotting was performed as described previously (11). Western blots from three independent biological replicates were quantified using the software ImageJ, version 1.0.

### Immunofluorescence microscopy

HCTT116 wt and NASP-dTAG cells were seeded at a density of 3000 cells per well in 96 well plates (6055300, Cell carrier) and treated with either dTAG-13 (5 μM, 48 hours) or DMSO (0.2%, 48 hours). Before fixation, newly replicated DNA was labelled with EdU (20 mins, 10 μM) at 37 °C. Cells were washed in ice-cold PBS, fixed (4 % paraformaldehyde, 15 mins, 4 °C) and washes twice in ice cold PBS. EdU staining followed the Click-iT plus Alexa647-picolyl azide protocol (Thermo Scientific) and DNA was stained with DAPI (4’,6-diamidino-2-phenylindole). Images were acquired on an Olympus ScanR high-content microscope and analysed with ScanR analysis software. Cell cycle gates were defined using mean EdU and total DAPI intensities.

### Data visualisation

Scatter and bar plots were visualised in GraphPad Prism (v9). All statistical analysis was performed in GraphPad Prism and statistical tests are detailed in figure legends.

## RESULTS

### Structural basis of sNASP dimerization

The human NASP has two splicing variants, sNASP and tNASP, with tNASP containing a longer acidic region (Figure 1A; ref 12,13). We have focused on structural studies of the TPR domain, responsible for sNASP and tNASP recognition of histones H3 and H4, and for the nucleosome assembly activity (31,36,45). To facilitate crystallization, we made a construct containing the TPR domain and capping helix region of human sNASP with the acidic region deleted (a.a 30–340 with a deletion Δ101–159; hereafter referred to as ‘sNASP core’, sNASPc). During gel-filtration purification, sNASPc was shown to have two peaks (conformations) on the chromatogram. We collected fractions from each peak and re-run the gel-filtration assays; both conformations, which were expected to be dimer and monomer, appeared to be highly stable without obvious exchange (Supplementary Figure S1A). The dimer and monomer conformations were further corroborated by the assays of size-exclusion chromatography coupled to multi-angle light scattering (SEC-MALS) (Figure 1B and Supplementary Table S1). Interestingly, the human sNASP had been shown to homodimerize *in vitro* (15,33).

**Figure 1.**
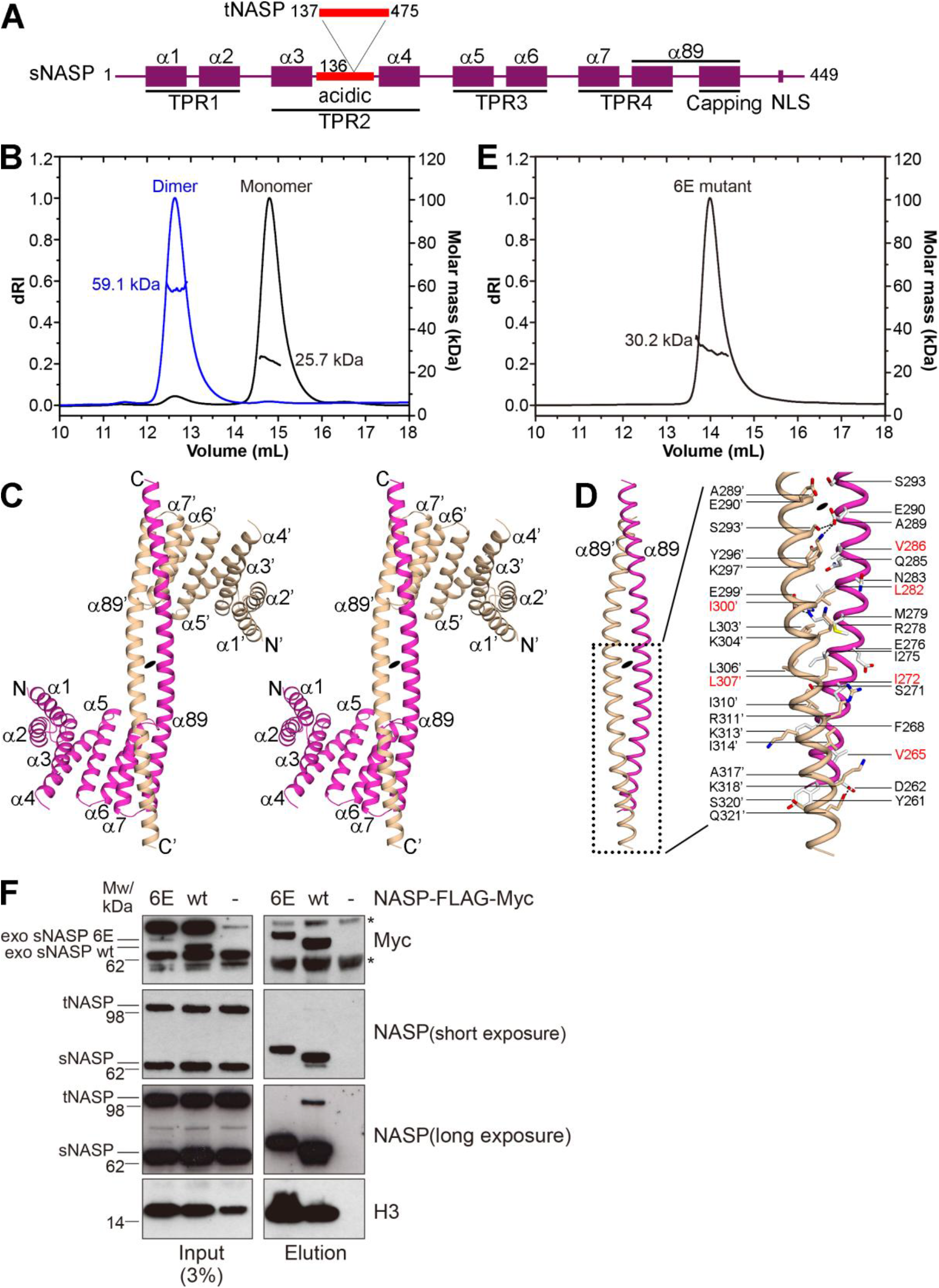
Structural and biochemical analysis of the sNASPc dimer. (**A**) Schematics of domain architectures of human sNASP and tNASP. tNASP has a longer acidic region with a 339-redidue segment inserted after position 136 of sNASP. (**B**) SEC-MALS analysis of sNASPc dimer and monomer. (**C**) Wall-eyed stereoview of ribbon representation of the structure of the sNASPc domain-swapping dimer. The two protomers, sNASPc and sNASPc’, are colored with magenta and wheat, respectively. (**D**) Cartoon view of the antiparallel coiled-coil like structure consisting of α89 and α89’. Highlighted the interactions between α89 and α89’. (**E**) SEC-MALS analysis of sNASPc 6E mutant. The residues of 6E mutant are highlighted in red in panel D. (**F**) Immunoprecipitation of sNASP-FLAG-Myc from HeLa S3 cells transiently transfected with wt and 6E mutant sNASP constructs or untransfected control cells (-). *, unspecific band.

After extensive efforts, we obtained crystals from the dimeric gel-filtration fraction and solved a 3.30 Å crystal structure of sNASPc in dimer conformation (stereo view in Figure 1C and Table 1). The two protomers in the structure are related by a 2-fold symmetric axis with each protomer containing four TPR motifs. Each of the TPR1-3 motifs adopts a canonical fold, whereas the TPR4 is a variant with the predicted second helix of TPR4 and the capping region together forming a very long and extended α-helix (namely α89 here). The extended α89 is constituted of residues from Tyr261 to Gly324 (64-residues long) of sNASPc with a kink at Glu305, and the kinked α89 helix is totally about 95 Å (65 Å + 30 Å) in length (Supplementary Figure S1B). The α89 and α89’ twist one another in a left-handed manner to form an antiparallel coiled-coil like structure, with the C-terminal one third of α89’ of the sNASPc’ protomer further packing against α7 and the N-terminal one third of α89 of the sNASPc protomer, and vice versa (Supplementary Figure S1C). Thus, the C-terminal one third of the α89 and α89’ helices each function like a ‘capping helix’, forming the unique architecture of a domain-swapping and dumbbell-shape dimer (Figure 1C).

**Table 1.** Data collection and refinement statistics

| | sNASPc dimer<br>(PDB 7V1K) | sNASPc–H3 α3<br>complex<br>(PDB 7V1L) | sNASPc–H3–H4–<br>ASF1b heterotetramer<br>(PDB 7V1M) |
| --- | --- | --- | --- |
| **Data collection** |  |  |  |
| Space group | P6<sub>2</sub>22 | C222<sub>1</sub> | P1 |
| Cell dimensions |  |  |  |
| *a*, *b*, *c* (Å) | 95.36, 95.36, 206.23 | 66.90, 177.05, 65.89 | 71.95, 71.90, 93.43 |
| $\alpha$, $\beta$, $\gamma$ (°) | 90, 90, 120 | 90, 90, 90 | 70.67, 70.64, 83.61 |
| Resolution (Å) | 40.0–3.30<br>(3.42–3.30)<sup>a</sup> | 40.0–2.85<br>(2.95–2.85) | 40.0–2.85<br>(2.95–2.85) |
| *R*<sub>pim</sub> (%) | 3.5 (42.0) | 4.5 (39.0) | 6.0 (47.1) |
| *I*/$\sigma$(*I*) | 17.0 (1.0) | 12.7 (1.0) | 10.2 (1.0) |
| *CC*1/2 | 0.518<sup>b</sup> | 0.475 | 0.606 |
| Completeness (%) | 99.9 (99.8) | 99.8 (98.4) | 95.6 (91.3) |
| Redundancy | 33.2 (25.9) | 12.0 (10.0) | 3.6 (3.3) |
| **Refinement** |  |  |  |
| Resolution (Å) | 39.18–3.29 | 36.74–2.85 | 35.59–2.85 |
| No. reflections (unique) | 9,000 | 9,478 | 37,598 |
| *R*<sub>work</sub> / *R*<sub>free</sub> (%) | 23.9/26.1 | 20.1/24.6 | 19.2/22.9<sup>c</sup> |
| No. atoms |  |  |  |
| Protein | 1740 | 1743 | 8136 |
| Peptide |  | 109 |  |
| *B*–factors |  |  |  |
| Protein | 131.8 | 72.8 | 72.3 |
| Peptide |  | 73.8 |  |
| R.m.s. deviations |  |  |  |
| Bond lengths (Å) | 0.009 | 0.008 | 0.011 |
| Bond angles (°) | 0.876 | 0.935 | 1.101 |
| Ramachandran plot |  |  |  |
| Outliers (%) | 0.0 | 0.0 | 0.1 |
| Favored/allowed (%) | 97.7/2.3 | 97.4/2.6 | 95.6/4.3 |

<sup>a</sup> Values in parentheses are for the highest-resolution shell. One crystal was used for each data set.
<sup>b</sup> *CC*1/2 of the highest-resolution shell are shown.
<sup>c</sup> The dataset of 7V1M was twinned, and a merohedral twin law (-*k*, -*h*, -*l*) was applied only for the final round of refinement in PHENIX.

The coiled-coil like interactions between α89 and α89’ constitute of totally 66 residues (33 residues from each helix), of which the predominant interactions are hydrophobic interactions and van der Waals contacts, with a few hydrogen bonds and salt bridges further stabilizing the dimer (Figure 1D). Moreover, most of these interacting residues are conserved amongst different species (Supplementary Figure S2A). Consistent with the extensive interactions, many different mutants with only single or even double mutations along the dimer interface failed to disrupt the dimer, and only a mutant with substitutions of six hydrophobic residues by glutamate (V265E I272E L282E V286E I300E L307E; Hereafter referred to as 6E mutant) was found to abolish the dimer formation (Figure 1E).

In accordance with our structural observations, co-immunoprecipitation experiments from HeLa S3 showed that exogenous FLAG-myc-tagged sNASP wild type (wt) pulled down both endogenous s-and t-NASP, while the sNASP 6E mutant did not (Figure 1F). This supports that NASP can form both homo- and hetero- dimers *in vivo* via the α89-mediated coiled-coil like interactions. Notably, disrupting sNASP dimerization did not affect its histone binding *in vivo*, as both the wt and 6E mutant efficiently pulled down histone H3 (Figure 1F).

### Structural basis for sNASP recognition of H3

We next undertook structural and biochemical analysis of the interactions between sNASP and H3. As a first step, our isothermal titration calorimetry (ITC) assays revealed that both the α3 region (residues 116–135) and the N-terminal region (residues 1–59) of H3 bound to sNASPc with comparable affinities at micromolar scale (*K*d of 0.7 and 1.2 μM, respectively), and that the αN region (residues 40–59) of H3 constituted the major binding site of the N-terminal region (*K*d of 5.3 μM) (Figure 2A,B,E). Our ITC results further revealed that the acidic region of sNASP was not necessary for binding to either the αN and α3 regions of H3 (Supplementary Figure S3A,B). Previous work showed that the N-terminal 79 residues of H3 were sufficient for binding to sNASP and tNASP in cell extracts (19), whereas another report identified an interaction between sNASP and the C-terminal α3 helix of H3 (31). Our results thus nicely corroborate and reconcile these findings.

**Figure 2.**
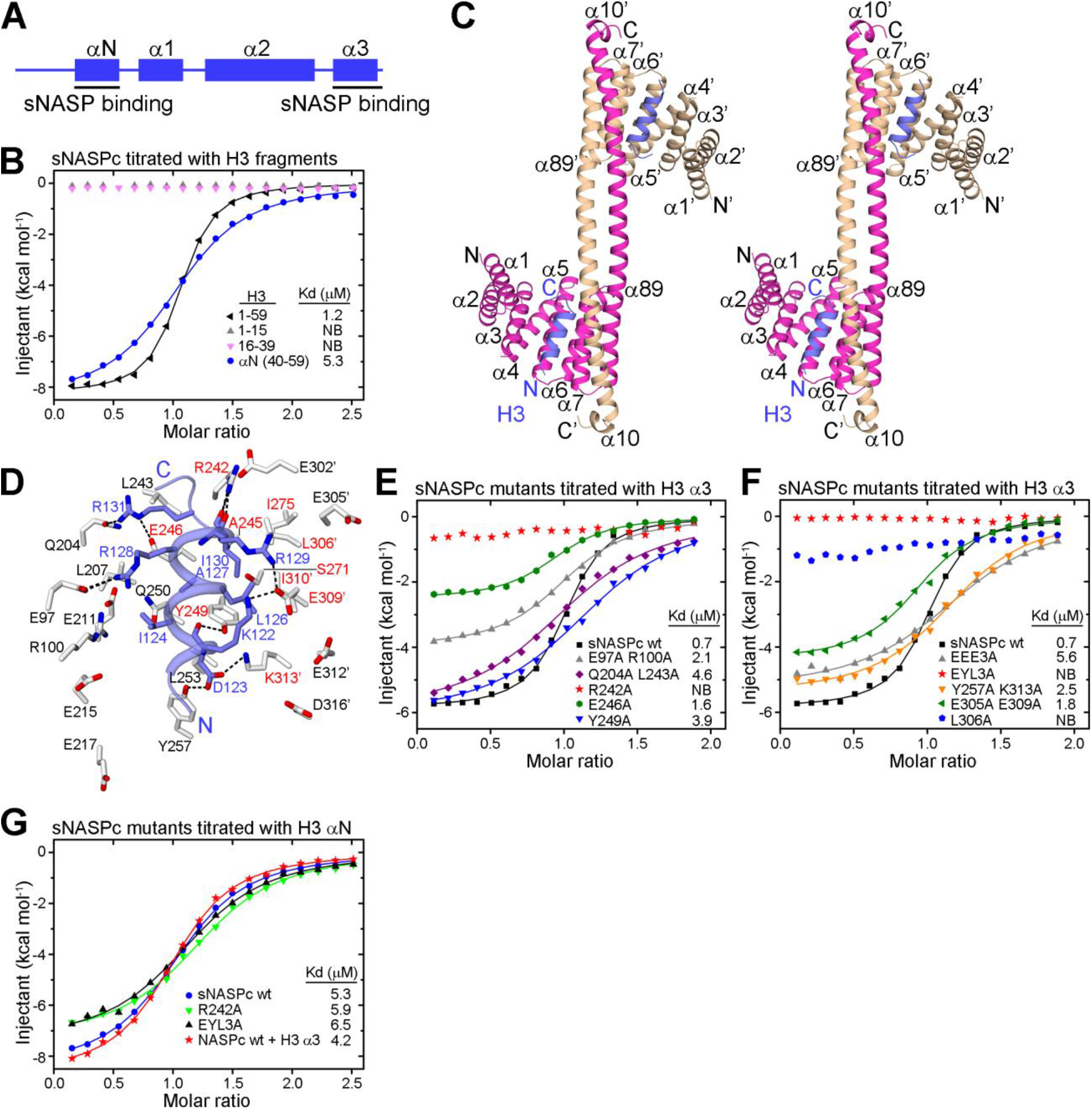
Structural and biochemical analysis of the interactions between sNASPc and H3. (**A**) Schematics of domain architecture of H3. (**B**) ITC analysis of sNASPc titrated with H3 N-terminal fragments. (**C**) Wall-eyed stereoview of ribbon representation of the structure of the sNASPc dimer in complex with the α3 region of H3. The two protomers of sNASPc and sNASPc’ are colored in magenta and wheat, respectively, whereas the two molecules of H3 α3 are colored in blue. (**D**) Zoom in view on the α3-binding site of the sNASPc–H3 α3 structure. Interacting residues of sNASPc (white) with H3 α3 (blue) are shown in sticks representation. Hydrogen bonds are indicated by dashed blacklines. The residues consisting of the hydrophobic pocket are labeled in red. (**E-F**) ITC analysis of sNASPc mutants in the α3-binding groove, titrated with H3 α3 peptide. (**G**) ITC analysis of sNASPc mutants in the α3-binding groove and sNASPc pre-bound with H3 α3, titrated with H3 αN peptide.

To obtain further insights into the recognition of H3 by sNASP, we solved a 2.85 Å crystal structure of the sNASPc dimer in complex with the α3 region of H3 (stereo view in Figure 2C). Each of the two H3 α3 molecules adopt a helical conformation highly similar to the one in the nucleosome structure and are encompassed within a groove in the concave side of each sNASPc protomer. The binding groove is formed by the conserved residues from the TPR3 and TPR4 motifs of one protomer and the ‘capping helix’ region of the other protomer, implying that dimerization is required for this binding mode. Moreover, the groove is amphipathic with a hydrophobic pocket located on one side and is surrounded by lots of conserved and negatively-charged residues (Supplementary Figure S3C,D).

The H3 α3 forms extensive interactions with the conserved residues of sNASPc that line up the groove (Figure 2D and Supplementary Figure S2A). The most prominent feature is that the main-chain carbonyl groups of H3 Arg129 and Ile130 establish three hydrogen bonds with the side chain of sNASPc Arg242, thus forming a helix-capping interactions. Another feature is that the side chain of H3 Leu126 and Ile130 are bound into the hydrophobic pocket formed by Arg242, Ala245, Glu246 and Tyr249 in NASPc α7, as well as Ser271 and Ile275 in sNASPc α8, and Leu306’, Glu309’, Ile310’ and Lys313’ from sNASPc’ α89’. It appears that the binding of H3 α3 can stabilize the sNASPc:sNASPc’ dimer. Moreover, five of the charged residues from H3 α3 contribute several salt-bridge and hydrogen-bonded interactions: both Lys122 and Arg129 of H3 establish salt bridges with Glu309’ of sNASPc’ α89’; Asp123 forms two hydrogen bonds to Tyr 249 and Tyr257 in sNASPc α7 and a salt bridge to Lys313’ in sNASPc’ α89’; Arg128 of H3 has two salt bridges with Glu97 of sNASPc α3; and Arg131 of H3 has a salt bridge and a hydrogen bond with Glu246 in sNASPc α7 and Gln204 in sNASPc α5, respectively. The van der Waals contacts among Ile124 and Ala127 of H3 with some surrounding sNASPc residues may further stabilize the interactions between H3 α3 and sNASPc. It should be noted that some of the negatively-charged residues lined up the groove, including Glu215 and Glu217 of sNASPc and Glu302’, Glu305’, Glu312’ and Asp316’ of sNASPc’, are not directly involved in binding to H3 α3.

Our ITC assays revealed that the sNASPc single mutants R242A and L306A and the triple mutant E246A Y249A L253A (EYL3A), all disrupted complex formation between sNASPc and H3 α3, whereas other sNASPc mutants reduced the binding by 2–8 folds (Figure 2E,F). These are consistent with our structural observations that Arg242 of sNASPc is involved in the prominent helix-capping interactions; Glu246, Tyr249 and Leu253 of NASPc are located at the center of the groove with Glu246 and Tyr249 further included in forming the hydrophobic pocket; Leu306’ of sNASPc’ is also involved in forming the hydrophobic pocket (Figure 2D). Interestingly, Glu246, Tyr249 and Leu253 were previously identified to be important for the interaction between sNASP and H3 α3 (31). Our ITC results showed that the αN region of H3 bound to sNASPc and mutants R242A and EYL3A, as well as to sNASPc pre-bound with H3 α3 with similar affinities (Figure 2G). This revealed that the H3 α3-binding groove on sNASPc was not involved in binding to the αN region of H3 and that sNASPc can simultaneously bind to the αN and α3 regions of H3. Taken together, our structure and biochemical analysis provide the molecular basis of how sNASP engages H3.

### Structural basis for the co-chaperone relationship of sNASP and ASF1b

sNASP is a major histone chaperone for H3–H4 dimers that cooperates with the histone chaperone ASF1 and the HAT1–RbAp46 acetyltransferase holoenzyme to maintain a soluble source of H3–H4 within the cell (15–19,46). The interactions among sNASP, H3–H4 dimers and ASF1 (a and b) had been studied through biochemical analysis (35), but the structural basis for this important and enigmatic co-chaperone complex is still elusive. To explore this, we mixed purified sNASPc dimers with H3–H4 tetramers and ASF1a (a.a. 1–155) at a molar ratio of 1:1:2 and successfully reconstituted a stable co-chaperone complex, the sNASPc–H3–H4–ASF1a heterotetramer. The stoichiometry of this complex was 1:1:1:1 as confirmed by the SEC-MALS analysis (Figure 3A). This observation suggests that sNASPc adopts a monomer conformation within the sNASPc–H3–H4–ASF1a heterotetramer, and that sNASPc transits from dimer to monomer conformation upon binding to the H3–H4 dimer and ASF1a. As our extensive attempts to crystallize the sNASPc–H3–H4–ASF1a heterotetramer were not successful, we constructed a covalent cassette of sNASPc-GGGSGGGS-ASF1b (a.a. 1–158) (referred to as sNASPc-8G-ASF1b) to facilitate crystallization, following similar designs used by us for structure determinations of several other co-chaperone complexes (10,38). Though with a covalent linker, the sNASPc-8G-ASF1b cassette formed a 1:1:1 complex with H3 and H4 in solution (hereafter referred to as the sNASPc–H3–H4–ASF1b heterotetramer; Figure 3A).

**Figure 3.**
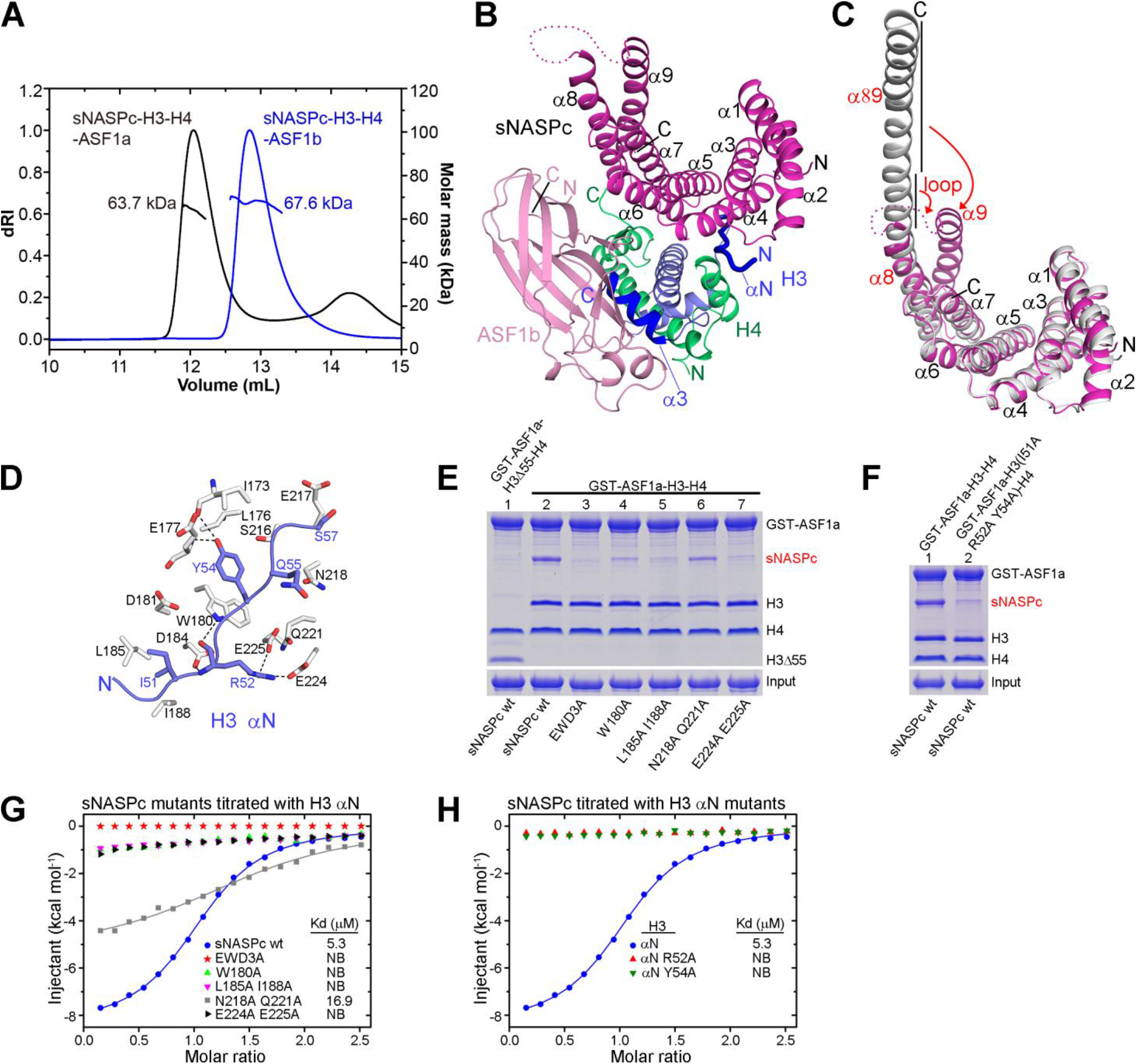
Structural and biochemical analysis of the sNASPc and ASF1b co-chaperone complex. (**A**) SEC-MALS analysis of the sNASPc–H3–H4–ASF1a complex and sNASPc–H3–H4–ASF1b complex. (**B**) Ribbon representation of the structure of the sNASPc–H3–H4–ASF1b complex. sNASPc, ASF1b, H3 and H4 are colored with magenta, pink, blue and green, respectively. The H3 αN and α3 regions are highlighted in dark blue. (**C**) Structural comparisons of the sNASPc monomer of the sNASPc–H3–H4–ASF1b structure with one of the protomer of the sNASPc dimer structure. Highlighting the conformational changes. (**D**) Zoom in view on the αN-binding site of the sNASPc–H3–H4–ASF1b structure. Interacting residues of sNASPc (white) with H3 αN (blue) are shown in sticks representation. Hydrogen bonds are indicated by dashed blacklines. (**E**) Pulldowns of sNASPc wt and mutants in the αN-binding site using GST-ASF1a–H3–H4 or GST-ASF1a–H3Δ55-H4. H3Δ55 indicates H3 with a deletion of the first 55 residues. (**F**) Pulldowns of sNASPc wt using GST-ASF1a–H3–H4 or GST-ASF1a–H3 (I51A R52A Y54A)–H4. The protein levels of sNASPc in panels E and F are quantified in supplementary Figure S4A and S4B, respectively. (**G**) ITC analysis of sNASPc mutants in the αN-binding groove, titrated with H3 αN peptide. (**H**) ITC analysis of sNASPc titrated with H3 αN peptides containing different mutation.

We solved a 2.85 Å crystal structure of the sNASPc–H3–H4–ASF1b heterotetramer (Figure 3B), in which sNASPc indeed adopted a monomer conformation. Comparisons of the sNASPc monomer and dimer structures reveal that the central 3 turns (residues 285–295) of the long α89 helix in the dimer structure had transited into a disordered loop in the monomer structure (Figure 3C). As a result, α89 separated into two helices, α8 and α9, in the monomer structure with α9 being folded back as a canonical capping helix. The TPR1-4 motifs in the monomer structure fit well with their equivalent parts in the dimer structure, and the α9 capping helix in the monomer structure also superimposes well with the C-terminal region of the α89’ helix in the dimer structure, with an overall rmsd of 0.77 Å (Supplementary Figure S3E). Interestingly, the structure of the H3–H4–ASF1b part within the sNASPc–H3–H4–ASF1b heterotetramer is highly similar to the reported structures of the ASF1–H3–H4 heterotrimer (Supplementary Figure S3F; ref 47,48). The α3 helix of H3 is sequestered and bound by ASF1b, thus the H3 α3-binding groove on sNASPc is unoccupied (Figure 3B), consistent with observations in a previous biochemical study (31). Moreover, sNASPc binds only to the αN region of H3 and, as expected (15,18,35), the interaction between sNASPc and ASF1b is bridged by the H3-H4 dimer. Notably, the αN region of H3 adopts an extended loop conformation, also observed when H3 is bound by factors that modify or bind K56 (49,50), but distinct from the α-helix it forms within the nucleosome structure (51).

The extended αN region of H3 interacts with several conserved residues from the α4 and α6 helices and the loop connecting the α5 and α6 helices (Loop56) on the convex surface of sNASPc (Figure 3D and Supplementary Figure S2A). The side chain of H3 Ile51 has lots of contacts with Asp181, Asp184, Leu185 and Ile188 in sNASPc α4. The main chain amide and carbonyl of H3 Arg52 are hydrogen bonded to the side chains of Trp180 and Asp184 in sNASPc α4, respectively, while its side chain forms salt bridges with Glu224 and Glu225 in sNASPc α6 and also has some contacts with Gln221 in sNASPc α6. The side chain of H3 Tyr54 penetrates into a pocket surrounded by Ile173, Leu176, Glu177 and Trp180 in NASPc α4 and Ser216 in sNASPc Loop56, with its aromatic ring establishing prominent stacking interactions with the side chain of sNASPc Trp180, and its hydroxyl group forming hydrogen bonds with the main chains of sNASPc Ile173 and Glu177. Furthermore, H3 Gln55 and Ser57 have some contacts with Asn218 and Glu217 in sNASPc Loop56, respectively.

To test the role of conserved residues for the sNASPc-H3 interactions within the sNASPc–H3–H4– ASF1b heterotetramer, we used the GST-ASF1a–H3–H4 complex to pull down sNASPc wt and mutants (lanes 2-7 in Figure 3E and Supplementary Figure S4A). As compared to the binding of sNASPc wt, the sNASPc mutants E177A W180A D181A (EWD3A), W180A, L185A I188A, N218A Q221A and E224A E225A reduced the binding to 5.5 ± 1.2%, 26.2 ± 5.5%, 13.6 ± 3.3%, 60.6 ± 6.6% and 11.1 ± 1.5%, respectively. Concurrently, H3 with a deletion in the αN region and the H3 mutant I51A R52A Y54A reduced the binding to 6.6 ± 1.5% (lane 1 in Figure 3E and Supplementary Figure S4A) and 16.0 ± 2.3% (Figure 3F and Supplementary Figure S4B), respectively. Moreover, our ITC assays with the same set of mutants showed that all of them, except the mutant sNASPc N218A Q221A, disrupted the formation of a complex between sNASPc and a H3 αN peptide (Figure 3G,H). Thus, these results support our structural observations and corroborate the importance of the sNASPc-H3 αN interactions for the integrity of the sNASPc–H3–H4–ASF1b co-chaperone complex.

### A model for engagement of H3–H4 dimers by sNASP

sNASP may engage H3–H4 dimers during early H3–H4 folding, and during transport, modification and storage of histones (4,19,26). After characterizing how sNASP recognize H3 and together with ASF1b co-chaperone a H3–H4 dimer; we further explored to how the dual H3 αN- and α3-binding modes of sNASP contribute to chaperoning H3–H4 dimers.

As compared to sNASPc wt, our pulldowns revealed that the sNASPc mutants EWD3A (H3 αN-binding deficient; Figure 3G) and EYL3A (H3 α3-binding deficient; Figure 2F) reduced the binding of H3–H4 dimer to 42.2 ± 2.7% and 52.0 ± 8.4% (quantification on H3), respectively, whereas the combinational mutant EWD3A+EYL3A further reduced the binding of H3–H4 dimer to 7.3 ± 3.4% (Figure 4A and Supplementary Figure S4C). Concurrently, the H3–H4 mutant with triple mutations I51A, R52A and Y54A in H3 (deficient in binding to the αN-binding site on sNASPc; Figure 3H) reduces the interaction of H3–H4 dimer with sNASPc wt to 57.2 ± 0.9%, and further reduces the interaction of H3–H4 dimer with sNASPc EYL3A to 9.5 ± 1.6% (Figure 4B and Supplementary Figure S4D). Similarly, pulldowns of sNASP^(30-340)^, a construct containing also the acidic region within TPR2, further support that the H3 αN- and α3-binding sites on sNASP are the major interaction sties for H3– H4 dimer *in vitro* (Supplementary Figure S4E,F). Interestingly, superimposition of the canonical H3– H4 structure/fold onto the H3 α3 helix bound by sNASPc shows lots of steric hindrances (Figure 4C), which suggests that the H3 α3 helix is disengaged from the histone fold to form the sNASP–H3–H4 complex. We thus speculated that a sNASP dimer can engage two partially unfolded H3–H4 dimers; the SEC-MALS assays of the sNASPc–H3–H4 complex did show a 2:2:2 complex and some higher-order complexes (Supplementary Figure S4G), which is consistent with previous observations (15). Moreover, pulldowns also reveal that the H3 αN- and α3-binding sites are conserved in frog (*X. laevis*) N1/N2, a homologue of sNASP implicated in the storage of H3–H4 dimers in Xenopus oocytes (26) (Figure 4D and Supplementary Figure S4H). Interestingly, though the structures of sNASPc and budding yeast homologue Hif1p (30) look similar with an overall rmsd of 2.2 Å (Supplementary Figure S5A), the dual H3 αN- and α3-binding modes were not conserved in Hif1p (Supplementary Figure S5B-D), as previously suggested for the α3-binding site (31). Taken together, these results thus suggest a model where sNASP by simultaneously holding the αN and α3 regions of H3 might engage a partially unfolded H3–H4 dimer *in vitro*, and the dual binding modes might also be implicated in H3–H4 storage by the frog N1/N2 protein.

**Figure 4.**
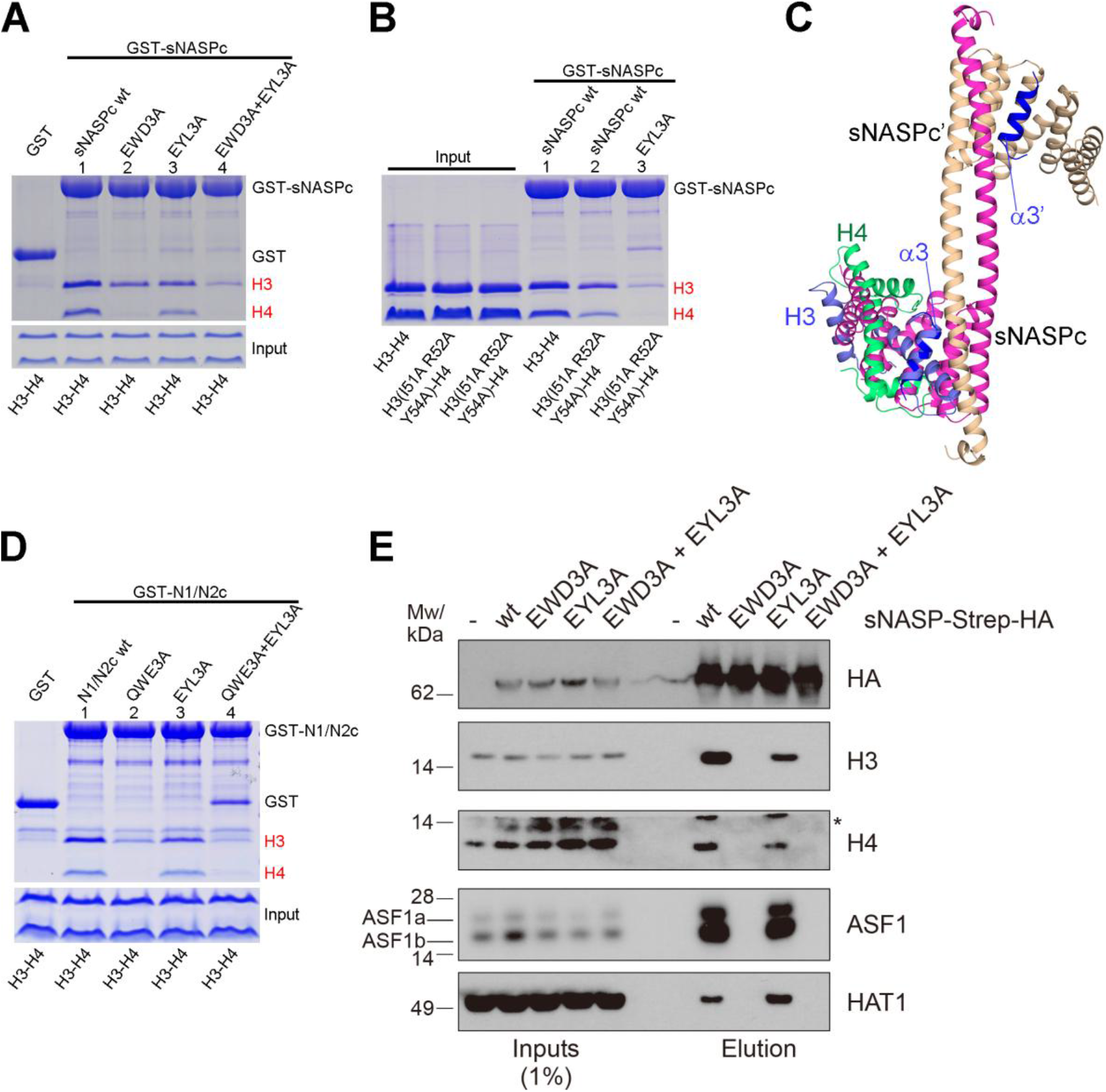
Interactions between sNASPc and H3–H4 dimers. (**A-B**) Pulldowns of histones H3–H4 by GST-sNASPc wt and mutants. The protein levels of H3 in panels A and B are quantified in Supplementary Figure S4C and S4D, respectively. (**C**) Superimposition of an H3–H4 dimer from the nucleosome structure (PDB 1KX5) onto our sNASPc–H3 α3 structure. The protomers NASPc and NASPc’ and the two molecules of H3 α3 are colored with magenta, wheat and dark blue respectively; the H3 and H4 from the H3–H4 dimer are colored with blue and green, respectively. Highlighting the steric hindrance between structures. (**D**) Pulldowns of histones H3–H4 by GST-N1/N2c wt and mutants. N1/N2c is ‘N1/N2 core’, containing residues 23–495 with the acidic region (a.a. 97–314) deleted. The protein levels of H3 are quantified in Supplementary Figure S4H. (**E**) Pulldown of Strep-HA-tagged sNASP from HeLa S3 cells induced to expressed the indicated mutants or uninduced control cells (-).*, unspecific band.

To test the importance of the dual H3 αN- and α3-binding modes for the chaperone function of sNASP *in vivo*, we performed a co-immunoprecipitation analysis of sNASP complexes from HeLa S3 cells conditionally expressing sNASP-Strep-HA wt and mutants (Figure 4E). We focused on ASF1 (a and b) and the histone acetyltransferase HAT1, both of which forms co-chaperone complexes with sNASP through mutual binding to histones (15,18). Compared to sNASP wt, the EWD3A mutant lost binding to both histones H3 and H4, ASF1 (a and b) and HAT1 (Figure 4E). In contrast, the EYL3A mutant only reduced H3 and H4 binding moderately but did not abrogate co-chaperoning with ASF1 or HAT1 binding (Figure 4E). These results validate the two independent H3 binding modes identified by our structural analysis and demonstrate that the interaction with the H3 αN region represents the major H3–H4 binding mode for sNASP in cells, and the one adopted in co-chaperone complexes with ASF1 (a and b) and HAT1. While the interaction of sNASP with the H3 α3 region is incompatible with mutual binding of ASF1 (a and b) to this site, this interaction could play a role during histone handover events where ASF1 (a and b) might transiently engage H3–H4 dimer through interaction with the H4 C-terminus.

Taken together, our results confirm the importance, both *in vitro* and *in vivo*, of sNASP dual binding to H3 αN and α3, suggesting that they both concurrently play a role in chaperoning H3 and H4 in the nucleosome assembly chain.

### Functional significance of NASP H3 αN- and α3-binding in vivo

NASP is essential for maintaining a soluble pool of histones H3–H4 during the cell cycle (19). To investigate the role of the H3 αN- and α3-binding modes in this function, we generated human HCT116 cells conditional for NASP expression by integrating a degradation tag (dTAG, FKBP12^F36V^) (52) in frame with both NASP alleles (Supplementary Figure S6A). In this background, we then conditionally expressed sNASP-Strep-HA wt and histone binding mutants. Addition of the dTAG-13 molecule allowed efficient and stable depletion of endogenous s- and t-NASP isoforms, demonstrated by western blotting of whole cell extracts (Figure 5A) and immunofluorescence (Supplementary Figure S6B). In accordance with previous results (19), loss of NASP reduced the pool of soluble histone H3–H4 (Figure 5B,C) without dramatic effects on cell cycle progression and cell proliferation (Supplementary Figure S6C,D). To measure the soluble histone pool, we quantified histone H3 and H4 levels separately as well as acetylation of H4 at lysine 5 (H4K5ac), a modification introduced by HAT1 on newly synthesized histones. Conditional expression of sNASP wt rescued the soluble histone pool in NASP-dTAG cells treated with dTAG-13 molecule (Figure 5C,D), indicating that tNASP is not specifically required for maintaining the pool. The H3 αN-binding mutant (EWD3A) failed to rescue the levels of H3, H4 and H4K5ac in NASP-dTAG cells (Figure 5C,D), demonstrating that binding to the H3 αN region is essential for sNASP chaperone function to maintain the soluble histone H3–H4 pool. Intriguingly, sNASP EYL3A efficiently rescued the levels of H4 and H4K5ac, but only partially restored H3 levels. Together and in agreeing with our structural and biochemical results, these data indicate that the H3 αN-binding mode is that primarily used by sNASP in chaperoning the H3–H4 histone pool, while the H3 α3-binding mode has a more specific role in regulating H3 metabolism.

**Figure 5.**
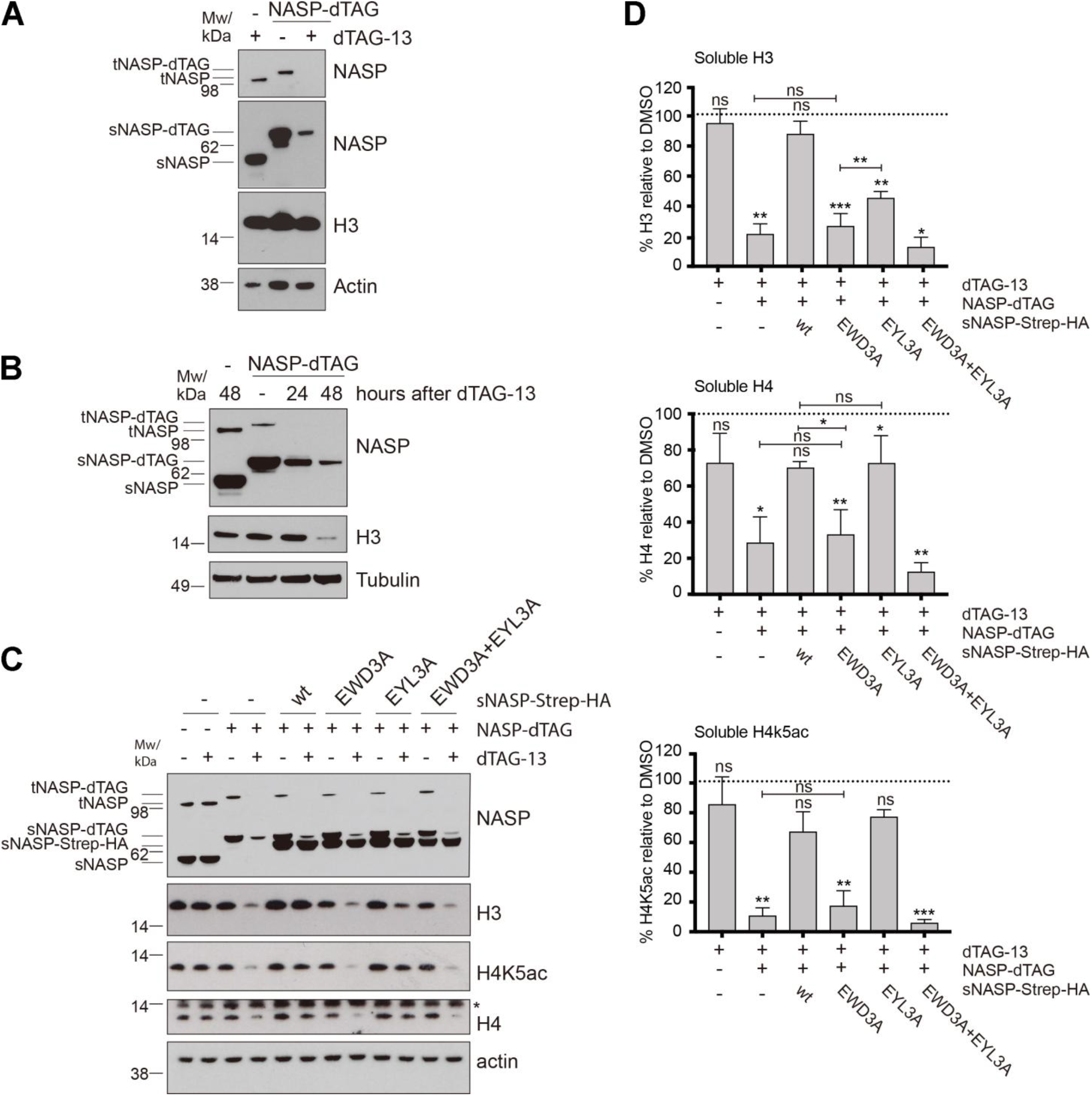
NASP maintains the soluble histone pool by chaperoning the H3 αN. (**A**) Western blot analysis of NASP in whole cell extracts of HCTT16 cells engineered to express NASP-FKBP12^F36V^ (NASP-dTAG) or wt control cells (-). Cells were treated with DMSO (-) or dTAG-13 for 48 hours as indicated. (**B**) Western blot analysis of soluble extracts from HCT116 NASP-dTAG and control cells (-) treated as described in panel A. (**C**) Western blot analysis of soluble extracts from HCT116 control cells (-) or NASP-dTAG (+) cells complemented as indicated with inducible expression of sNASP-Strep-HA wt or histone binding mutants. The expression of sNASP-Strep-HA was induced by DOX treatment for 48 hours and cells were co-treated with dTAG-13 for 48 hours as indicated. *, unspecific band. (**D**) Quantification of H3, H4 and H4K5ac from the Western Blot analysis in panel C. The mean is shown with s.d (n=3). Bands were quantified and normalized to actin and shown relative to DMSO treatment. P values represent unpaired two-sided t-tests (from left to right H3 quantification: P=0.8544; 0.0095; 0.5687; 0.0010; 0.7766; 0.0019; 0.0064; 0.0138; H4 quantification: P= 0.3438; 0.0348; 0.2500; 0.0069; 0.0427; 0.6578; 0.0387; 0.1262; 0.0016; H4K5ac quantification: P= 0.8761; 0.0017; 0.0720;; 0.0023; 0.6336; 0.1359; 0.0001).

## DISCUSSION

In conclusion, through structural and functional characterizations of sNASP and its interactions with histone substrates, we revealed the molecular basis of NASP as a major H3–H4 chaperone. Taken together this work supports that NASP operates through the H3 αN-binding mode to chaperone H3– H4 together with ASF1 and maintain the soluble histone H3–H4 supply. In addition, we propose that NASP, acting mainly upstream in the histone supply chain, chaperones monomeric H3 and partially folded H3–H4 using its H3 α3-binding mode.

Our structures were based on the construct sNASPc, containing the core of sNASP. As tNASP is only different from sNASP with an insertion within the acidic region (Figure 1A), the structures of sNASPc are therefore likely representative of both sNASP and tNASP. We also noted there is no specific requirement for tNASP in H3–H4 chaperoning in cancer cells (our work and Cook et al (ref 19)), consistent with most somatic cells lacking this isoform. Except being rich in D/E residues, the acidic region is without any conserved motif amongst species. We found that NASP used dual binding modes for interaction with histones H3 and H4, while the acidic region of NASP was not necessary for this interaction. Interestingly, the acidic region of NASP was reported to be necessary for the interaction with linker histone H1 (45), but the structural basis of how NASP chaperones H1 are still unknown.

NASP was reported to form both a monomer and a dimer *in vitro* (33). Our sNASPc structure showed that NASP forms a domain-swapping dimer through the long α89 helix. The residues involved in dimerization are mostly conserved, at least, amongst vertebrate species. Consistently, we observed that the recombinant frog N1/N2 can form both a monomer and a dimer in solution (Supplementary Figure S4I). We also found that sNASP could homodimerize, and even heterodimerize with tNASP, in human cells. The functional significance of NASP dimerization *in vivo* is still unknown. In the scenario of histone folding, one protomer of tNASP dimer might hold histones H3 and/or partially folded H3–H4 and the other protomer might bind to HSP90 as a co-chaperone, which thus promotes the folding (and dimerization) of an H3–H4 dimer. We also elucidated a NASP ‘monomer’ structure, obtained in the context of the sNASPc–H3–H4–ASF1b co-chaperone complex. The NASP dimer and monomer transition involved the transformation of the long α89 helix into the α8 helix and the α9 capping helix. We envision this transition between dimer and monomer may be responsive for NASP interactions with diverse histone substrates in different cellular context.

Our studies revealed the dual H3 α3- and αN- binding modes of NASP, the former previously suggested by NMR analysis (31). The structural and biochemical results showed that both the H3 α3- and αN- binding modes of NASP were involved in binding to H3–H4 dimers and that the H3 αN-binding mode of NASP was implicated in formation of the co-chaperone complex with ASF1 (a and b) *in vitro*. Interestingly, the cellular data showed that the H3 αN-binding mode was the major H3–H4 binding mode for NASP, and the one adopted in co-chaperone complexes with ASF1 (a and b) and HAT1 *in vivo*. Consequently, the H3 α3-binding mode seems to have limited role in chaperoning the major H3–H4 pool *in vivo*. However, in the histone supply chain, NASP binding to the H3 α3 may be important in histone handover events between NASP and ASF1, where this H3 α3 helix could be transiently exchanged between the two chaperones. Moreover, the H3 α3-binding mode fits well with NASP function in early folding steps of the H3–H4 dimer, as NASP would hold the H3 α3 which might let the H3 α1-α2 region in a good configuration to pair with H4. Several proteins participate in the folding of histones H3 and H4, including the newly identified histone chaperone DNAJC9 (11) and the molecular chaperones HSC70 and HSP90 (15,46). Thus, these chaperones might compensate NASP function when it is deficient in binding to the H3 α3 region in our experiments. Interestingly, the H3 α3-binding deficient mutant, EYL3A, rescued H4/H4K5ac levels while only partially rescuing H3 levels in our complementation experiments. This represents novel evidence that a fraction of H3 in the soluble fraction exists in the form of monomer, as suggested by Apta-Smith et al (20). However, we found that the H3 αN-binding mode is that primarily used by NASP in maintaining the H3–H4 dimer pool in cells, implying that shielding of the H3 N-terminus (especially the αN region) by NASP is necessary for protecting histones from degradation. ASF1 function is not required for protecting the soluble histone pool (19), underscoring that that shielding the αN region rather than co-chaperoning with ASF1 is critical in preventing degradation. This suggests that the free histone H3 tail could represent a signal for degradation or induce instability. In absence of NASP, soluble H3 and H4 are degraded by chaperone-mediated autophagy (19). The soluble histone pool is mostly composed by newly synthetized histones, with only a little contribution from evicted or old histones (53), which carry different sets of PTMs. However, following DNA damage (54,55) and transcription (56,57), the proportion of evicted histones could increase locally and it is unclear if they are also chaperoned by NASP and re-incorporated into chromatin. It would be interesting to explore whether PTMs on the H3 tail would influence its interaction with NASP and consequently the stability of evicted H3–H4 dimers. For example, the phosphorylation of H3 Tyr54 (a residue conserved in higher eukaryotes) is expected to disrupt the αN-binding mode of NASP and might thus induce H3–H4 dimer degradation. Such investigation could provide insight into whether new and evicted old histones, carrying different H3 tail PTMs, are handled by distinct histone metabolic pathways.

## Supporting information

This PDF file includes: Supplementary Figures S1 to S6 Supplementary Tables S1

## AVAILABILITY

Coordinates of the X-ray crystal structures have been deposited in the RCSB PDB (www.rcsb.org) with the following accession numbers: 7V1K (the sNASPc dimer), 7V1L (the sNASPc–H3 α3 complex), and 7V1M (the sNASPc–H3–H4–ASF1b heterotetramer). Other data are available upon reasonable request.

## ACCESSION NUMBERS

Atomic coordinates and structure factors for the reported crystal structures have been deposited in the Protein Data bank under accession numbers 7V1K (the sNASPc dimer), 7V1L (the sNASPc–H3 α3 complex), and 7V1M (the sNASPc–H3–H4–ASF1b heterotetramer).

## SUPPLEMENTARY DATA

Supplementary Data are available at NAR online.

## ACKNOWLEDGEMENT

We thank Drs Baixing Wu, Yongrui Liu and Fudong Li for technical assistance. We thank the staffs from the BL17B1/BL18U1/BL19U1/BL19U2/BL01B beamlines of National Facility for Protein Science in Shanghai (NFPS) at Shanghai Synchrotron Radiation Facility, for assistance during data collection.

