## Supplementary material for "NASP maintains histone H3–H4 homeostasis through two distinct H3 binding modes": This PDF file includes: Supplementary Figures S1 to S6 Supplementary Tables S1

Supplementary Figures S1 to S6
Supplementary Tables S1

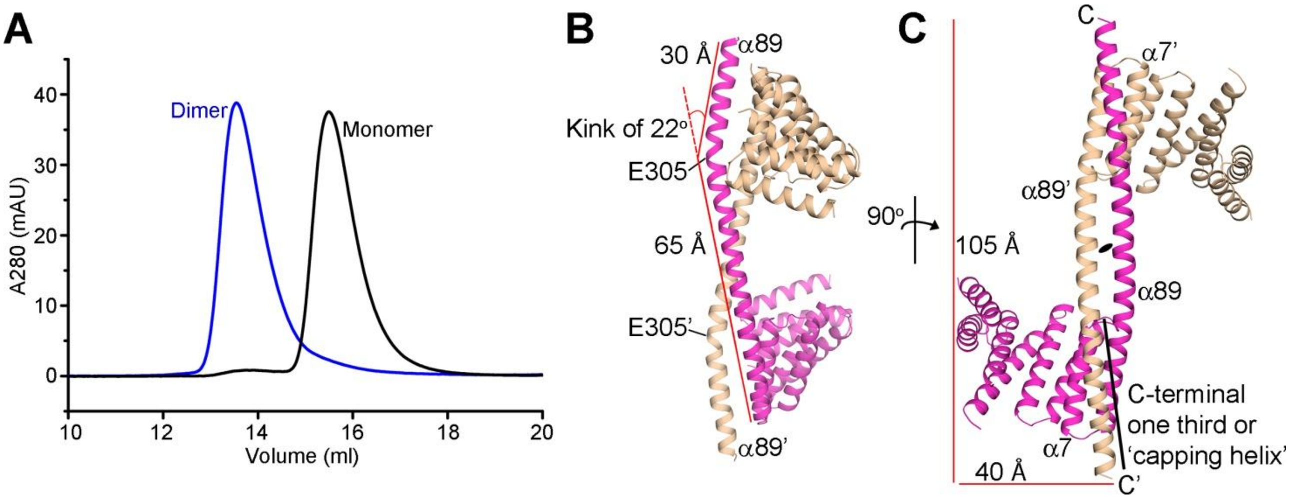


**Supplementary Figure S1.** Gel-filtration analysis and structure of sNASPc dimer. (**A**) Two peaks or conformations of sNASPc were found in the gel-filtration chromatogram during purification. The fractions from each peak were collected separately and re-loaded onto the Superdex 200 Increase 10/300 GL column (GE Healthcare) for analysis. The two conformations are very stable in our conditions and expected to be dimer and monomer. (**B-C**) Structures of the sNASPc dimer highlighting the Kink structures in the long helices α89 and α89' (**B**) and how the C-terminal one third of α89' of the sNASPc' protomer packs against α7 and the N-terminal one third of α89 of the sNASPc protomer, and vice versa (**C**). The two protomers sNASPc and sNASPc' are colored with magenta and wheat, respectively.

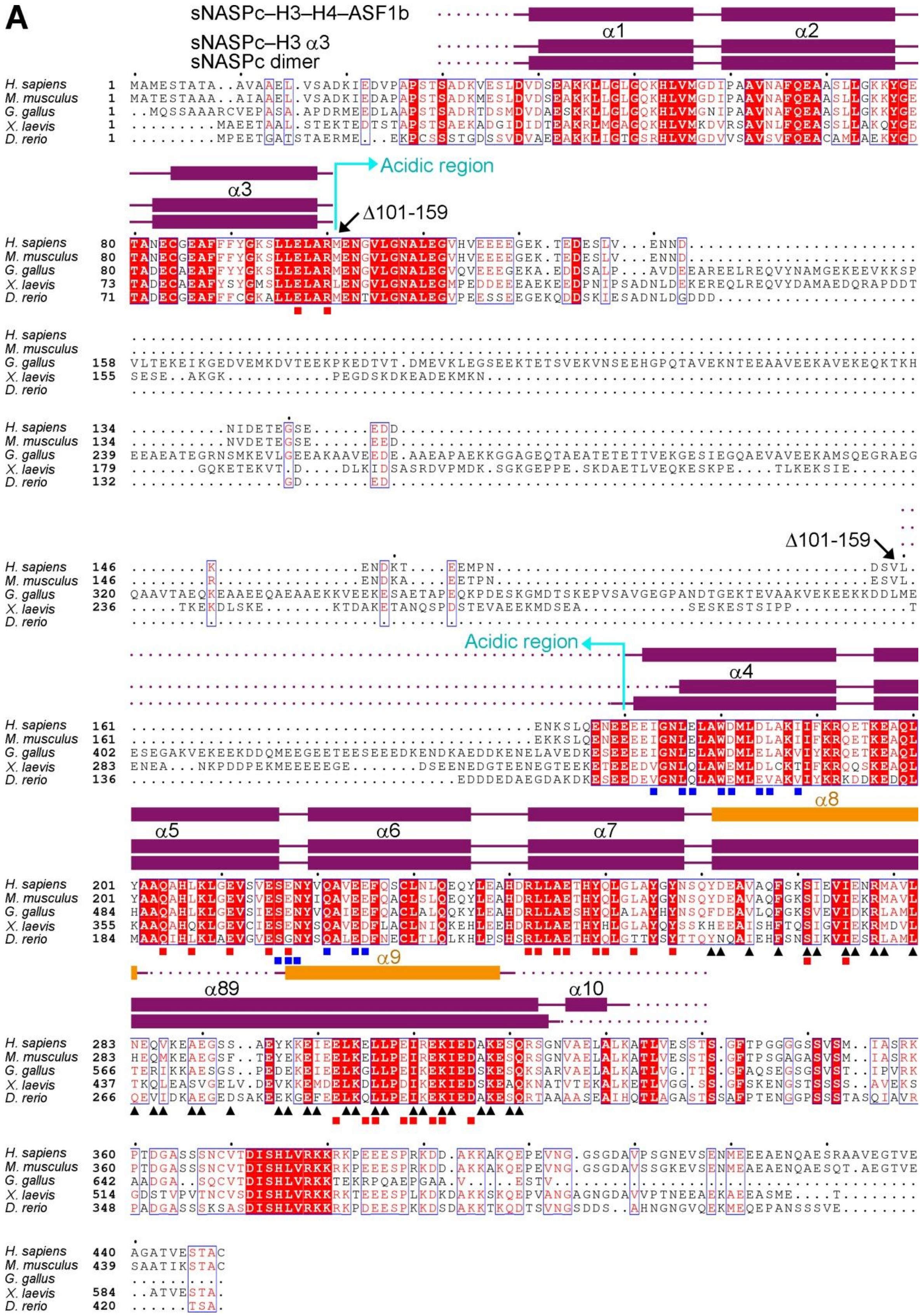
A
sNASPc–H3–H4–ASF1b
sNASPc–H3 α3
sNASPc dimer
α1
α2
α3
Acidic region
Δ101-159
α4
α5
α6
α7
α8
α9
α89
α10
H. sapiens
M. musculus
G. gallus
X. laevis
D. rerio

**Supplementary Figure S2.** Sequence alignment of NASP. (**A**) Sequence alignment of NASP from different species: *H. sapiens* sNASP (NP_689511); *M. musculus* sNASP (NP_001074944); *G. gallus* NASP (XP_015146756); *X. laevis* N1/N2 (NP_001081537); *D. rerio* sNASP (XP_021332627). The conserved and identical residues across species are boxed and highlighted in red. Secondary structure elements derived from the structures of the sNASPc dimer, sNASPc–H3 α3 complex and sNASPc–H3–H4–ASF1b heterotetramer are shown on top of the alignments. The disordered regions not resolved in the density maps of the crystal structures are indicated by magenta dots. Under the alignments, '▲' highlights the 33 residues in the α89 helix consisting of the dimerization interface of the sNASPc dimer; '■' highlights the 25 residues forming the H3 α3-binding groove of sNASPc; '■' highlights the 14 residues forming the H3 αN-binding site of sNASPc.

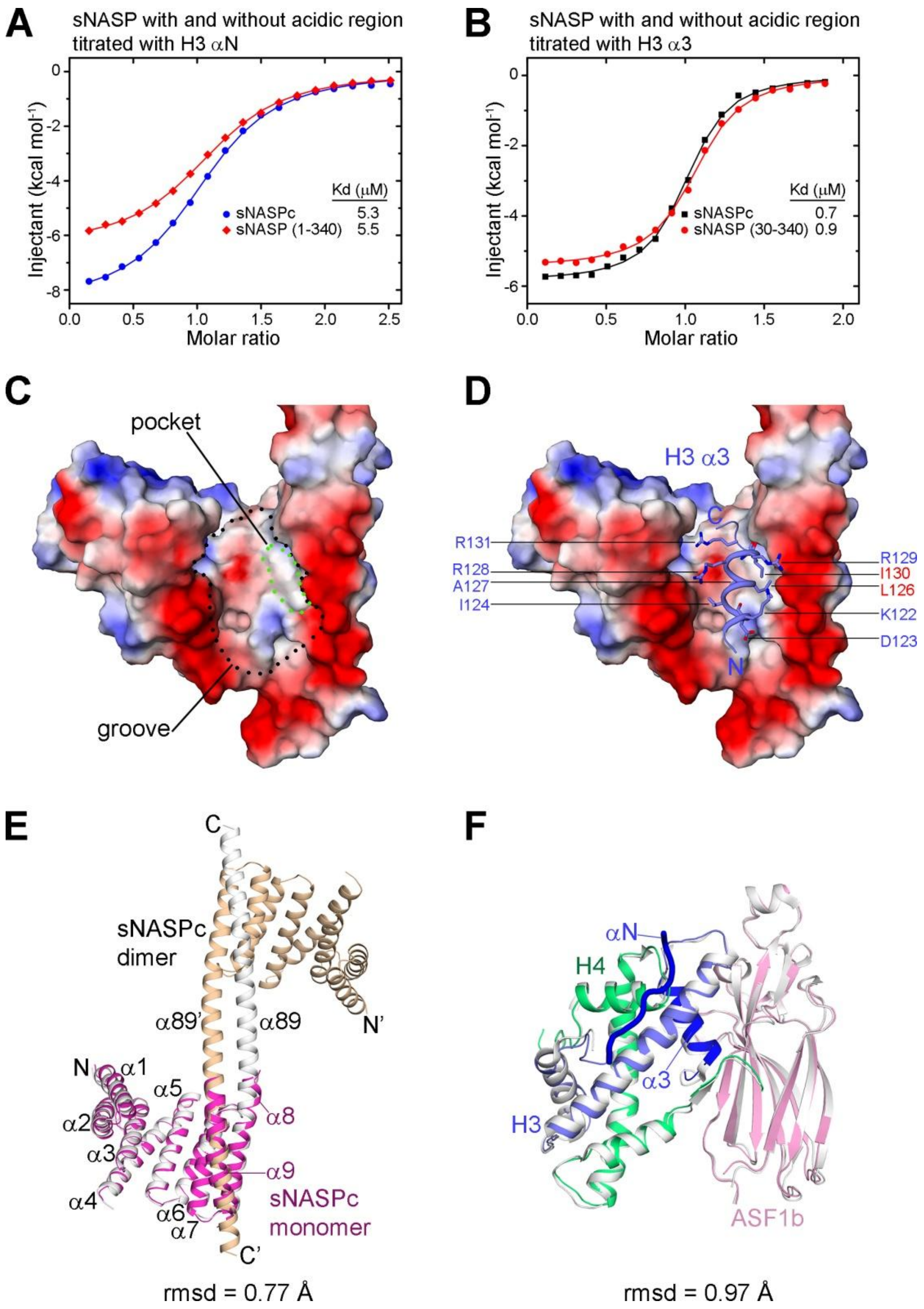


**Supplementary Figure S3.** Characteristics of the sNASPc–H3 α3 complex and the sNASPc–H3–H4–ASF1b heterotetramer. (**A-B**) ITC analysis of sNASP with and without the acidic region titrated with the H3 αN peptide (**A**) and the H3 α3 peptide (**B**), respectively. (**C-D**) Surface view of sNASPc color-coded with the electrostatic potential (red, negatively charged; blue, positively charged). The H3 α3-binding groove and the hydrophobic pocket within the groove are highlighted with black and green dashed circles, respectively (**C**). The H3 α3 bound in the groove is shown as ribbon representation (**D**); and the interacting residues of H3 α3 are shown in sticks representation (**D**). (**E**) Superimposition of the structure of the sNASPc monomer (colored in magenta) from the sNASPc–H3–H4–ASF1b heterotetramer onto the structure of the sNASPc dimer (two protomers in white and wheat, respectively). The rmsd of the two superimposed structures is 0.77 Å. (**F**) Superimposition of the structure of the H3–H4–ASF1b part (color coded as in Figure. 3B) from the sNASPc–H3–H4–ASF1b heterotetramer onto the known structure of the ASF1a–H3–H4 trimer (colored in white; PDB 2IO5). The H3 αN and α3 regions of the H3–H4–ASF1b part are highlighted in dark blue. The rmsd of the two superimposed structures is 0.97 Å.

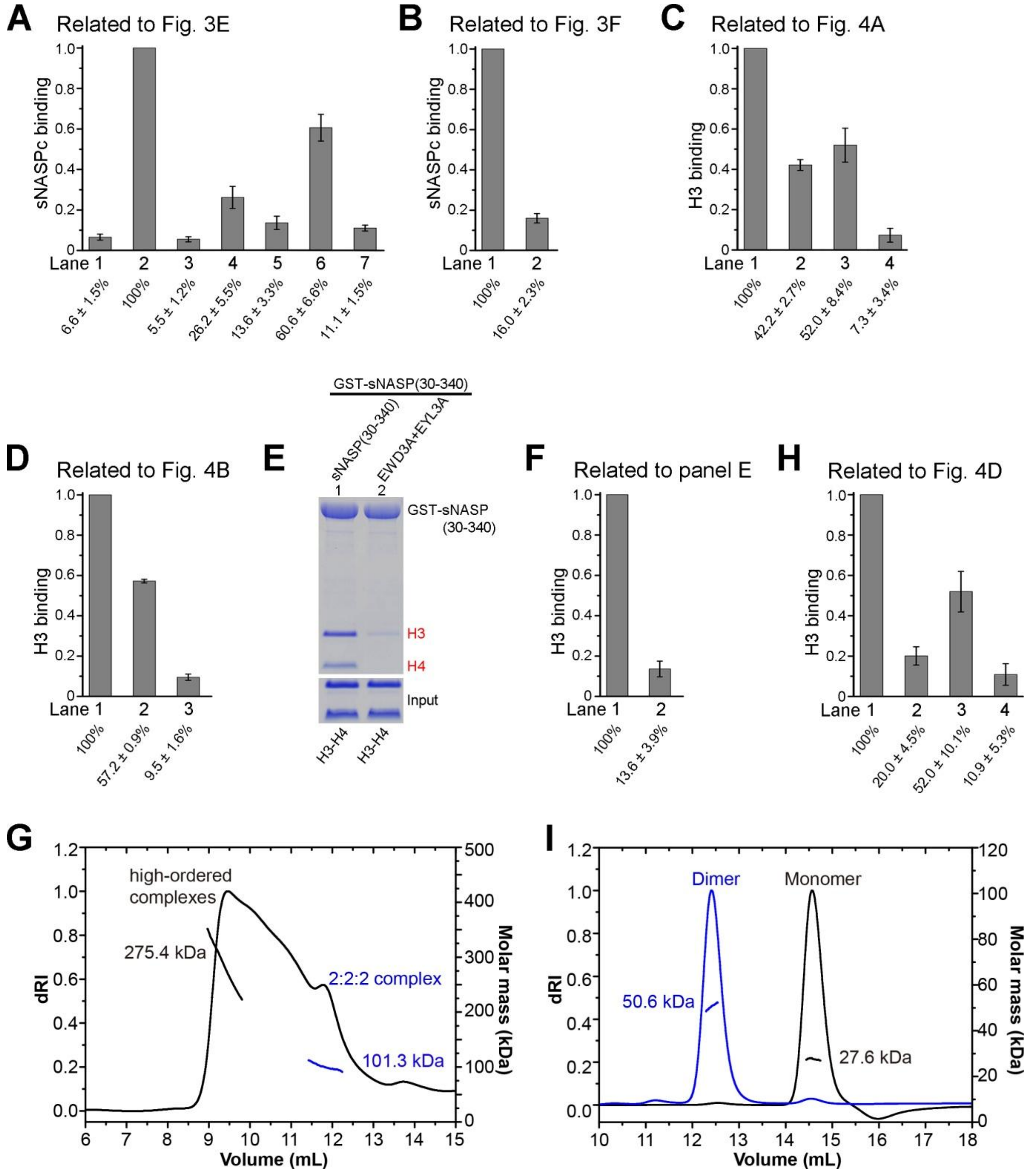


**Supplementary Figure S4.** Quantification of pulldown assays. (**A**) Quantification of pulldowns related to Figure 3E. The mean is shown with s.d (n=3). The levels of bound sNASPc wt and mutants are normalized with the level of sNASPc wt bound to GST-ASF1a–H3–H4 (lane 2) set as 100%. (**B**) Quantification of pulldowns related to Figure 3F. The mean is shown with s.d (n=3). The levels of bound sNASPc wt are normalized with the level of sNASPc wt bound to GST-ASF1–H3–H4 (lane 1) set as 100%. (**C**) Quantification of pulldowns related to Figure 4A. The mean is shown with s.d (n=3). The levels of bound H3 are normalized with the level of H3 bound to GST-sNASPc wt (lane 1) set as 100%. (**D**) Quantification of pulldowns related to Figure 4B. The mean is shown with s.d (n=3). The levels of bound H3 wt and mutants are normalized with the level of H3 wt bound to GST-sNASPc wt (lane 1) set as 100%. (**E**) Pulldowns of GST-sNASP$^{(30\text{-}340)}$ wt and mutant with histones H3–H4. The levels of H3 are quantified in panel F. (**F**) Quantification of pulldowns related to panel E. The mean is shown with s.d (n=3). The levels of bound H3 are normalized with the level of H3 bound to GST-sNASPc$^{(30\text{-}340)}$ wt (lane 1) set as 100%. (**G**) SEC-MALS analysis of the sNASPc–H3–H4 complex. (**H**) Quantification of pulldowns related to Figure 4D. The mean is shown with s.d (n=3). The levels of bound H3 are normalized with the level of H3 bound to GST-N1/N2c wt (lane 1) set as 100%. (**I**) SEC-MALS analysis of N1/N2c dimer and monomer.

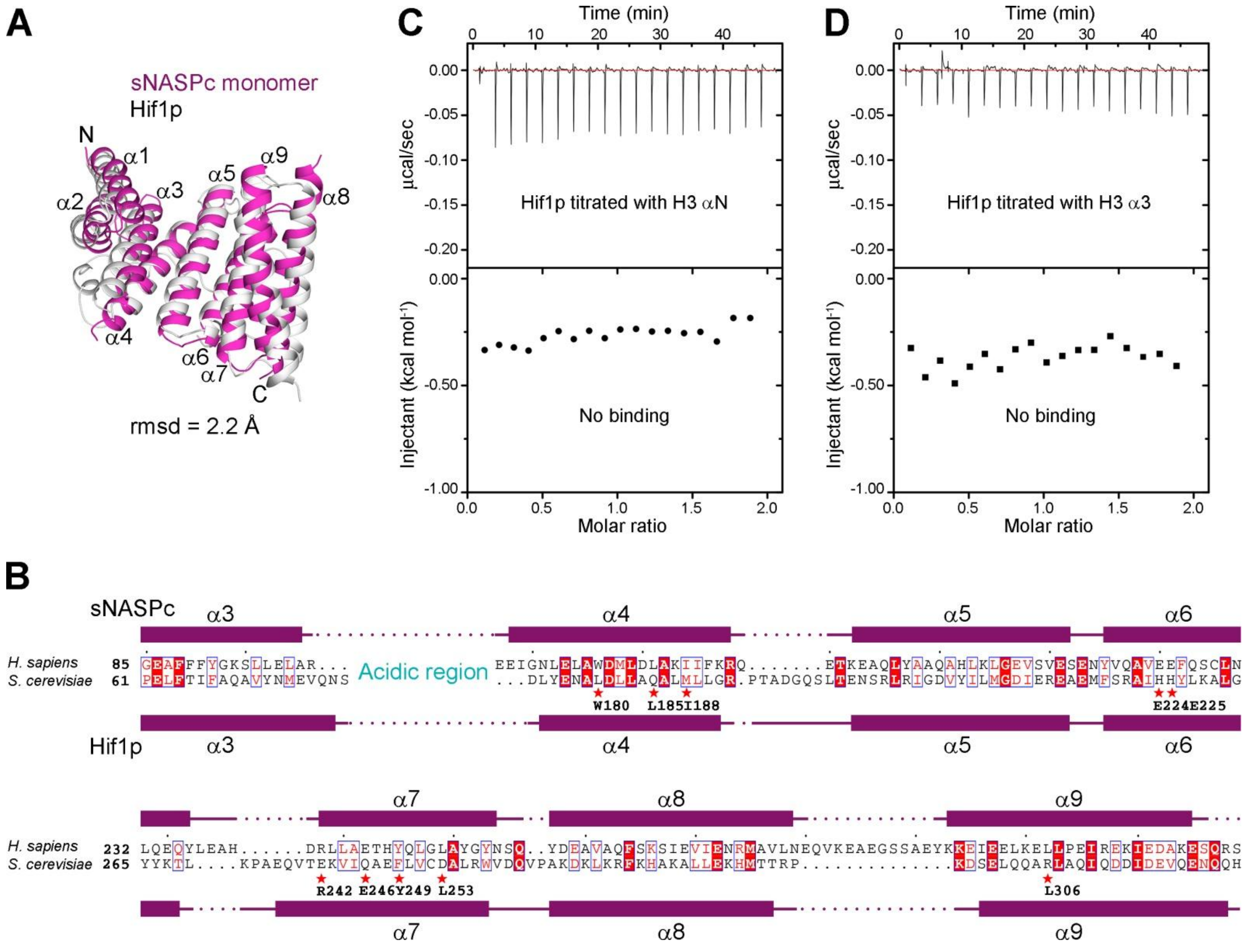


**Supplementary Figure S5.** Comparisons of sNASP and budding yeast Hif1p. (**A**) Superimposition of the structure of the sNASPc monomer (colored in magenta) derived from the sNASPc–H3–H4–ASF1b heterotetramer onto the structure of budding yeast Hif1p (colored in white; PDB 4NQ0). The superimposition covered the TPR2-4 motifs and the capping helices of sNASPc and Hif1p, whilst the TPR1 motifs did not fit well in the two structures. The rmsd of the two superimposed structures is 2.2 Å. (**B**) Sequence alignment of *H. sapiens* sNASP (NP_689511) and *S. cerevisae* Hif1p (NP_013078). The alignment was focus on the TPR2-4 motifs and the capping helices, whilst the TPR1 motifs and the acidic regions were not conserved and omitted for alignment. The alignment was manually adjusted based on structural superimposition in panel A. The conserved and identical residues are boxed and highlighted in red. Secondary structure elements derived from the structures of the sNASPc monomer and budding yeast Hif1p (PDB 4NQ0) are shown on top and bottom of the alignment, respectively. Under the alignments, '★' highlights the key residues of sNASP in the H3 αN-binding site (W180, L185, I188, E224 and E225) and H3 α3-binding site (R242, E246, Y249, L253 and L306), illustrating that some of these key residues are not conserved in Hif1p. (**C**) ITC analysis of Hif1p titrated with the H3 αN peptide, which did not show any obvious interaction. (**D**) ITC analysis of Hif1p titrated with the H3 α3 peptide, which did not show any obvious interaction. This confirmed the result from Bowman et al (1).

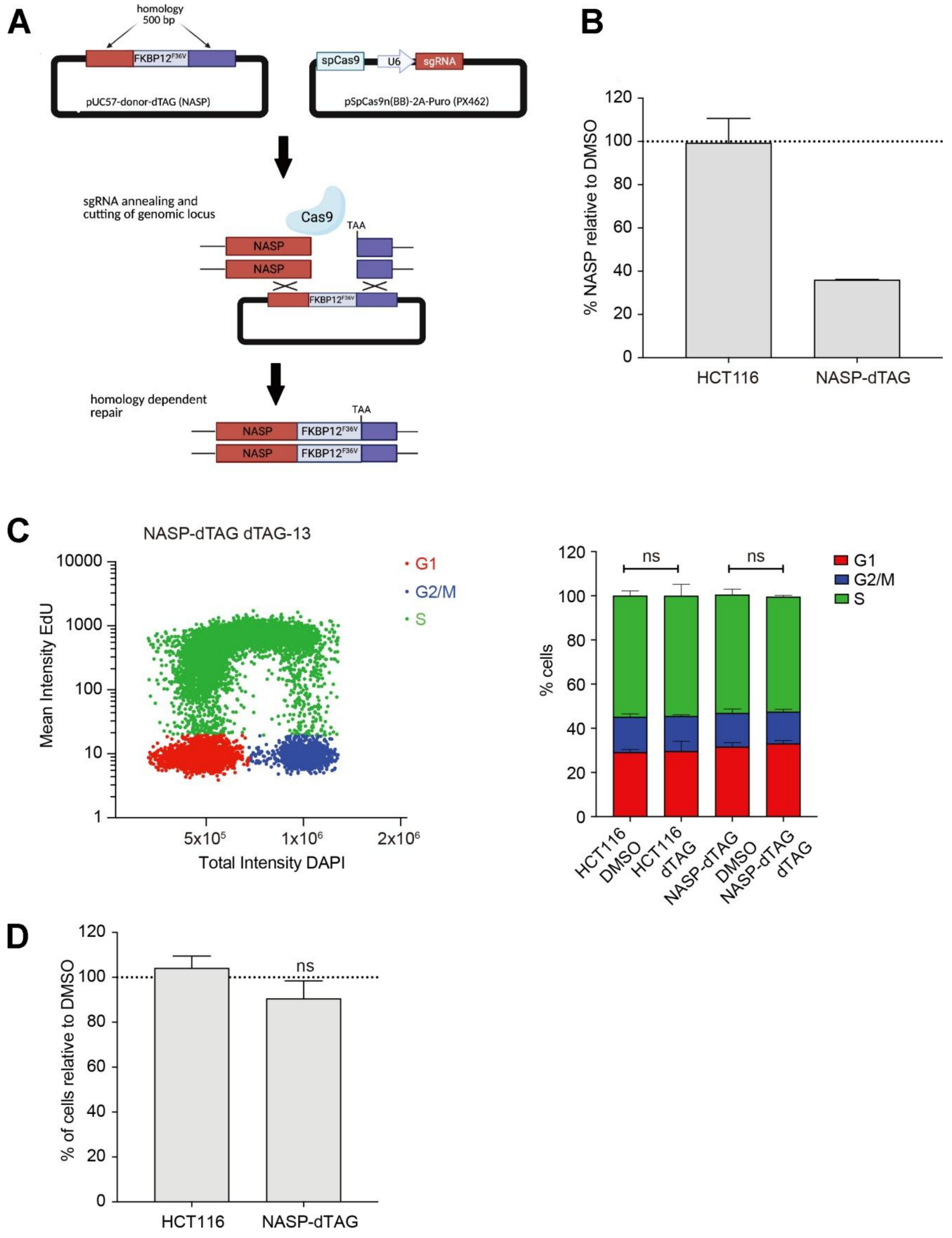


**Supplementary Figure S6.** Characterization of NASP-dTAG cell line. (**A**) Schematic depiction of the $FKBP12^{F36V}$ knock-in strategy. (**B**) High-content microscopy of HCT116 wt and NASP-dTAG cells treated with DMSO or dTAG-13 for 48 hours and pulsed with EdU before fixation. The NASP signal was measured in the nucleus and shown relative to the DMSO control. Bars represent the mean with s.d. (n=3). (**C**) Cell cycle distribution measured by high-content microscopy in cells treated as in panel b. (left panel) Representative diagram illustrating gating strategy for quantification of G1, S and G2/M populations. (right panel) Bar-diagram showing cell cycle distribution across HCT116 wt and NASP-dTAG cells treated as indicated. Mean is shown with ± s.d (n=3). ns, non-significant indicates P>0.05 in multiple unpaired t test (from left, G1 P= 0.873142; 0.349609; S phase P=0.919112; 0.550069; G2 P=0.908600; 0.366155). (**D**) Cell viability measured by cell titer blue in HCT116 wt and NASP-dTAG cells treated with DMSO or dTAG-13 for 48 hours. Viability is shown relative to the DMSO control with bars indicating the mean with s.d. (n=3). ns, non-significant indicates P>0.05 in multiple unpaired t test (from left to right, P= 0.837880; 0. 0.311381)

**Supplementary Table S1.** Results of SEC-MALS assays.

| Complex | Expected stoichiometry | Expected mass (kDa) | Measured mass* (kDa) | Measured mass/Expected mass |
|---|---|---|---|---|
| sNASPc (dimer peak) | 2 | 56.4 | 59.1 | 1.05 |
| sNASPc (monomer peak) | 1 | 28.2 | 25.7 | 0.92 |
| sNASPc 6E mutant | 1 | 28.3 | 30.2 | 1.07 |
| sNASPc–H3–H4–ASF1a(1-155) | 1:1:1:1 | 72.1 | 63.7 | 0.88 |
| sNASPc-8G-ASF1b(1-158)–H3–H4 | 1:1:1 | 71.1 | 67.6 | 0.95 |
| sNASPc–H3–H4 | 2:2:2 (latter peak) | 109.2 | 101.3 | 0.93 |
| | 2×(2:2:2) or 3×(2:2:2) (former peak; higher-order complexes) | 218.4 or 327.6 | 275.4 | 1.26 or 0.84 |
| N1/N2c (dimer peak) | 2 | 57.2 | 50.6 | 0.88 |
| N1/N2c (monomer peak) | 1 | 28.6 | 27.6 | 0.97 |

* Buffer conditions: 20 mM Tris, pH 7.5, 500 mM NaCl.
